# A two-stage approach to identifying and validating modifiable factors for the prevention of depression

**DOI:** 10.1101/759753

**Authors:** Karmel W. Choi, Murray B. Stein, Kristen Nishimi, Tian Ge, Jonathan R.I. Coleman, Chia-Yen Chen, Andrew Ratanatharathorn, Amanda B. Zheutlin, Erin C. Dunn, 23andMe Research Team, Major Depressive Disorder Working Group of the Psychiatric Genomics Consortium, Gerome Breen, Karestan C. Koenen, Jordan W. Smoller

**Keywords:** Resilience, protective factors, polygenic risk, depression, prevention, PheWAS

## Abstract

**Background:** Although depression is recognized as the leading cause of disability worldwide, decades of research have identified few actionable preventive factors. Using phenotypic and genomic data from the UK Biobank, we took advantage of a unique opportunity to screen a wide range of potentially modifiable factors that could offset known risk factors for depression.

**Methods:** We curated baseline data on more than 100 lifestyle and environmental factors in participants’ lives, including behavioral (e.g., exercise, sleep, media use, diet), social (e.g., support, activities), and environmental (e.g., greenspace, pollution) variables. In a follow-up survey, participants reported on their traumatic life experiences and mental health, including depression. Polygenic risk scores for depression were generated based on large-scale genome-wide association results. Excluding those meeting criteria for depression at baseline, we identified at-risk individuals at high predicted probability (> 90th percentile) for clinically significant depression at follow-up based on their (i) polygenic risk, or (ii) reported traumatic life events. Using a factors-wide design corrected for multiple testing and adjusted for potential confounders, we identified modifiable factors associated with follow-up depression in the full sample and among at-risk individuals. Using a two-sample Mendelian randomization (MR) design, we then examined which significant factors showed potential causal influences on depression risk, or vice versa.

**Results:** A range of baseline modifiable factors were prospectively associated with follow-up depression, including factors related to social engagement, physical activity, media use, and diet. MR follow-up analyses provided further support for the effects of social support-seeking, TV use, and other factors on depression risk.

**Conclusion:** As the field increasingly quantifies the role of genetic factors in complex conditions such as depression, knowledge of modifiable factors that could offset one’s genetic risk has become highly relevant. Here, we present an approach to screening for potentially modifiable factors that may offset the risk of depression in general and among at-risk individuals. In light of the burden of disease associated with depression and the urgent need for actionable preventive strategies, this approach could help prioritize candidates for follow-up studies including clinical trials for depression prevention.

## Introduction

Depression now represents the leading cause of disability worldwide^1^, highlighting a pressing need for effective treatment and prevention strategies. However, despite decades of research into the causes and consequences of depression, our knowledge of actionable strategies, including modifiable factors that could mitigate depression risk at the population level, remains relatively limited. Two of the best substantiated risk factors for depression—genetic vulnerability and early life adversity^2,3^—are effectively unmodifiable in adults.

A number of critical research gaps are evident. First, the literature to date has primarily focused on validating a limited set of candidate modifiable factors such as physical activity^4,5^ or social support^6^. While theoretically grounded and informative, there may be additional factors that could modify depression risk but remain overlooked or unknown. Scanning a wider range of factors could help confirm existing relationships while potentially identifying novel targets for prevention strategies. Systematically testing the relationship between many variables and a single outcome for hypothesis-free discovery is now common practice in other fields in the form of genome-or phenome-wide association studies and has led to new insights about underlying associations^7,8^, but has not been applied to searching for modifiable factors for depression.

Second, few studies to our knowledge have assessed the relative influence of multiple modifiable factors in the same population. Although individual studies may pursue intriguing variables (e.g., dietary factors), the influence of these factors on depression—while statistically significant in a narrower context—may not be robust or as clinically important when considered with other factors. Understanding the relative importance of modifiable factors for depression has been limited to date by inadequate sample sizes for multiple testing and the lack of comprehensive measurements of modifiable factors in a single study, but is now enabled by large cohort studies such as the UK Biobank^9^ in which wide-ranging assessments are available.

Third, we do not always know which modifiable factors can make a difference for individuals who are at particularly elevated risk for genetic and/or environmental reasons. Some factors that generally help prevent depression in the general population may not necessarily be most relevant for individuals with specific risk profiles^10^, and vice versa. Genetic vulnerability represents an important source of risk for depression, with heritability estimates over 40%^2^.

Similar to many other psychiatric disorders, depression is now recognized as a polygenic condition^11^—influenced by many variants across the genome with individually small effects^12^. As we are increasingly able to quantify polygenic risk for depression^13^ and potentially return this information to individuals in the future^14^, it becomes vital to expand knowledge of effective actionable measures for those identified at elevated risk. In addition to genetics, life history factors such as traumatic events are known to increase risk for depression^15^. As we more comprehensively assess established sources of genetic and environmental risk in a precision medicine framework^16^, knowledge of modifiable factors that are most relevant for high-risk individuals could help guide recommendations on how to offset pre-existing vulnerabilities for depression^17^.

Finally, modifiable factors may be associated with depression in observational data for myriad reasons, including unaccounted third variables (i.e., confounding) as well as reverse causation (e.g., whereby depression risk influences observed behavioral patterns). To strengthen conclusions about which factors may be high-priority intervention targets, Mendelian randomization (MR) analyses can be used to further test the relationship between empirically identified modifiable factors and depression. MR is an alternative strategy for potential causal inference that uses genetic variants inherited at birth to approximate a natural experiment in which individuals are assigned to varying average lifetime levels of an exposure (e.g., social affiliation) in relation to an outcome of interest (e.g., depression)^18^. While MR also has its limitations, its use of genetic data bypasses typical sources of confounding in observational data and allows for independent triangulation of traditional questions^19^. We previously leveraged the MR framework to validate a relationship between objectively measured physical activity and reduced risk of depression^5^. Here, we extend this framework to examine a range of potential factors that may influence depression.

In the present study, we took advantage of a unique opportunity to screen a broad range of potentially modifiable factors for depression. Using phenotypic and genomic data from UK Biobank participants without substantial depressive symptoms at baseline, we conducted association tests between 105 potentially modifiable factors and clinically significant depression at follow-up (Figure 1). Given the established causal role of genetics and traumatic life events on depression risk, we also aimed to identify modifiable factors that may influence depression even in the context of these largely static risk factors. Finally, we used two-sample Mendelian randomization analyses to further assess the directional and potentially causal relationships between identified modifiable factors and depression.

**Figure 1.**
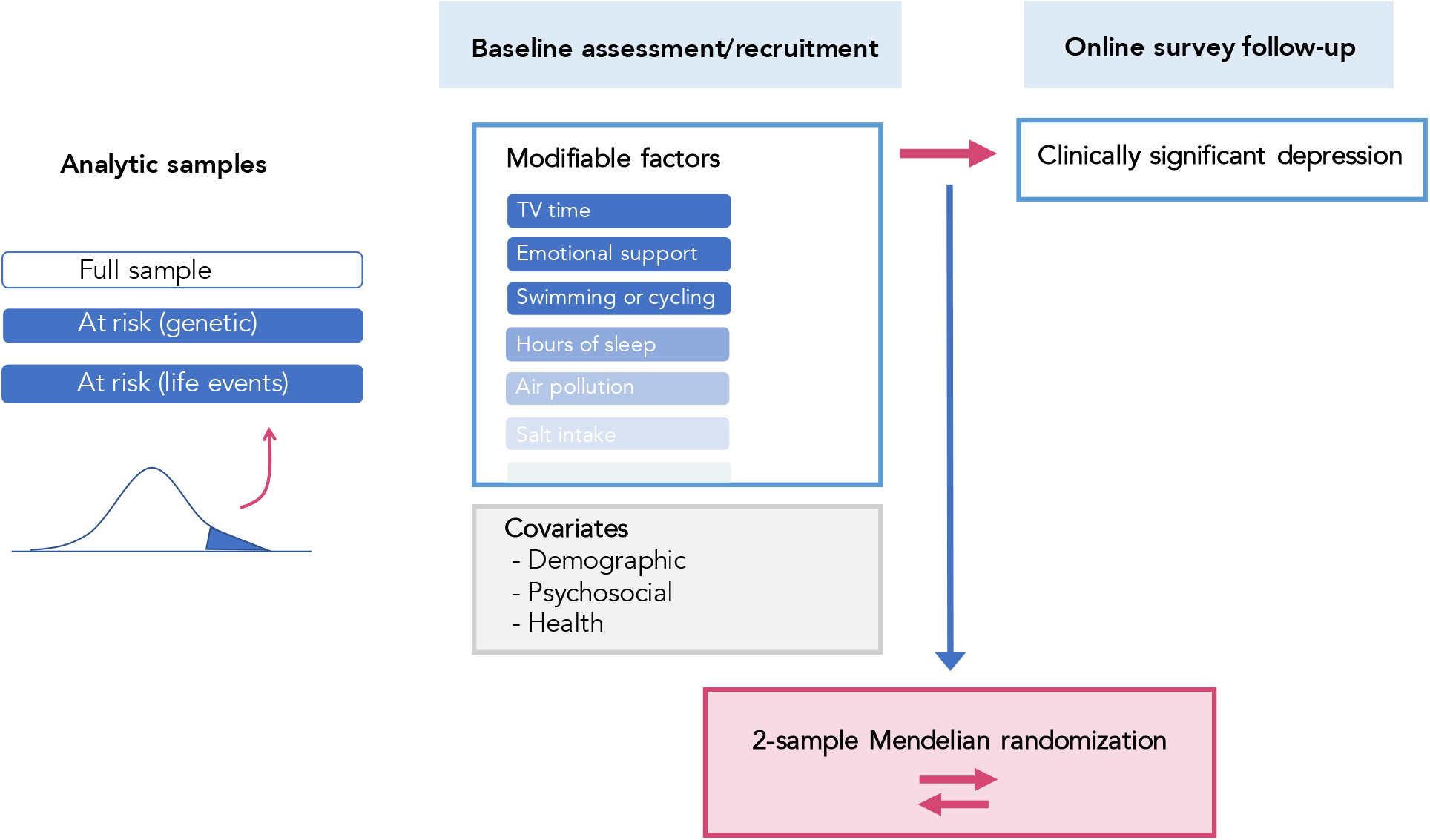
Overview of analytic design to test prospective associations between modifiable factors and subsequent depression. Associations were tested in three analytic samples: (a) full sample; (b) at-risk individuals based on polygenic risk; and (c) at-risk individuals based on reported traumatic life events. To reduce bias in associations from contemporaneous reporting, modifiable factors were selected from those indexed to the baseline assessment, while subsequent depression was assessed at the follow-up survey approximately 5 years later. Key distinctions with previous depression analyses in the UK Biobank are summarized in **Supplementary Methods S0**, emphasizing our targeted focus on modifiable factors for depression in a prospective design and among different risk groups.

## Methods

### Sample and Procedures

Our sample consisted of 123,794 adult individuals of white British ancestry who enrolled in the UK Biobank, had high-quality genomic data (for quality control procedures, see **Methods S1**), and completed an online follow-up mental health survey approximately five years after their initial enrollment (Figure 1). Data analytic procedures were approved by the institutional review board at Partners HealthCare and conducted as part of UK Biobank application #32568. Data processing and statistical analyses were conducted between October 2018 and July 2019.

### Measures

#### Depression

At baseline, participants reporting depressed mood and/or anhedonia (for details, see **Methods S2**) for more than half the days in the past two weeks were considered to have elevated depressive symptoms^20^ and were excluded (n=5,416; leaving n=118,378). Follow-up symptoms of depression were then measured in an online survey approximately five years after the baseline assessment, using the PHQ-9^21^, which were summed to create an overall score ranging from 0 to 27. To derive predicted probabilities for depression to stratify at-risk groups, we created a binary variable for clinically significant depression at follow-up based on a score cut-off of >10^22^.

#### Modifiable factors

We curated data on 105 potentially modifiable factors that were measured or derived at the baseline assessment (**Table S1**). These factors included behavioral (e.g., exercise, sleep, media use, diet), social (e.g., activities, support), and environmental (e.g., greenspace, pollution) variables. We selected these variables by inspecting the UK Biobank online data showcase (http://biobank.ctsu.ox.ac.uk/crystal/index.cgi) for broad domains of modifiable factors and relevant variables within these domains. After review by three separate authors (KWC, JWS, KN), we included variables in a domain that were (a) not likely a close comorbidity of mental health (e.g., substance use or cognitive functioning); (b) putatively modifiable at an individual and/or societal level (e.g., lifestyle or environmental factors); and (c) largely available for most participants and not just administered to a small subset (e.g., based on branching response options). Potentially correlated variables within a category (e.g., 16-hour and 24-hour noise pollution) were retained to assess the relative influences of all available variables. We also selected two non-modifiable variables hypothesized to be unrelated to depression, i.e., hair color and skin tanning tendency, as negative controls. Data cleaning and processing were performed on all variables (described in **Methods S3 and Table S1**). Continuous variables were scaled to a mean of 0 and standard deviation of 1.

#### Traumatic life experiences

Participants also reported on their history of traumatic life experiences in the online follow-up survey—including childhood physical, sexual, and emotional abuse; partner-based physical, sexual, and emotional abuse; and other lifetime traumatic events, specifically, sexual assault, violent crime, life-threatening accident, and witnessing violent death (for details, see **Methods S2**).

#### Covariates

Baseline variables were also extracted for basic participant characteristics (i.e., participant sex, age, assessment center); sociodemographic factors (i.e., socioeconomic deprivation, employment status, household income, completion of higher education, urbanicity, household size); and physical health factors (i.e., BMI, and physical illness/disability) (for details and inclusion rationale, see **Methods S2**).

### Polygenic risk scoring

Polygenic risk scores (PRS) were generated based on large-scale genome-wide association results for major depression^11^—specifically, we used summary statistics (discovery GWAS n=431,394) from the Psychiatric Genomics Consortium leaving out the UK Biobank to minimize sample overlap and including 23andMe data for improved statistical power. We retained SNPs with minor allele frequency > 0.01 and INFO quality score > 0.80 for scoring. To generate polygenic risk scores, we applied PRS-CS^23^—a Bayesian polygenic prediction method that places a continuous shrinkage (CS) prior on effect sizes for all HapMap3 SNPs and infers posterior SNP weights using GWAS summary statistics combined with an external LD reference panel (1000 Genomes Project European sample). Because PRS-CS (available as a Python package via https://github.com/getian107/PRScs) allows multivariate modeling of local LD patterns and can accommodate a range of underlying genetic architectures while preserving all SNPs for scoring, it demonstrates increased explanatory power compared to conventional and other Bayesian methods, particularly when using a large discovery GWAS^23^ (for comparison with conventional clumping and thresholding, see **Methods S4**). We set the global shrinkage parameter at 0.01 to reflect the likely polygenic architecture of major depression. Scores were calculated by summing the number of risk alleles at each SNP multiplied by the posterior SNP weight inferred using PRS-CS, with a total of 1,090,207 included SNPs. We standardized the scores within this analytic sample (mean=0, SD=1; for distribution, see **Methods S4**). We then extracted residuals from a model in which PRS were regressed against the top 10 European ancestry PCs provided by the UKB for use as stratification-adjusted PRS in subsequent analyses.

### Stratifying participants at risk for follow-up depression

Among individuals with available data on later depression and risk variables (i.e., polygenic risk and reported traumatic life events) (n=113,587; 5% meeting criteria for follow-up depression), we removed a holdout training sample of 1,000 participants consisting of an even split of randomly selected cases and controls for follow-up depression (for rationale, see **Methods S5**). In this holdout training sample, we regressed follow-up depression against (a) polygenic risk, or (b) reported traumatic life events. For the latter, each traumatic life event was entered as a separate independent variable within a multivariable model to estimate relative weights of each event on depression risk, rather than assuming equal influences. We obtained regression model coefficients for each set of risk variables from the training sample (**Methods S5**) and used these coefficients as weights to generate predicted probability scores for follow-up depression for individuals in the remaining testing sample (n=112,587)—based on (a) polygenic risk, or (b) reported traumatic life events (for resulting distributions, see **Methods S5**). Selecting individuals with high predicted probability scores (> 90th percentile) for depression, we obtained three samples: (i) individuals in the full sample unselected for risk (*full*; maximum n=112,587), (ii) individuals at higher risk based on genetic factors (*PRS*; maximum n=11,258), and (iii) individuals at higher risk based on reported traumatic life events (*TLE*; maximum n=11,258). Only 13.8% of participants (n=1,558) could be assigned to both PRS and TLE risk groups, suggesting only modest overlap and potentially distinct influences on depression (for exploratory results in this reduced sample, see **Figures S20-25)**.

### Factors-wide association scan

Using a factors-wide association approach with logistic regression (**Methods S5**), we tested associations between each baseline modifiable factor and clinically significant follow-up depression in each of these samples (Figure 1), with a conservative Bonferroni-corrected threshold for establishing top hits (p = 0.000159, i.e., 0.05 divided by 105 tests in three primary analytic samples). All associations were adjusted for participant sex, baseline age, and assessment center (Model 0). We further adjusted for potential sociodemographic confounders described earlier (Model 1), then added physical health factors (Model 2). All analytic samples were restricted to participants with full covariate data (*full*: maximum n=100,519; *PRS*: maximum n=10,093; *TLE*: maximum n=10,152) to ensure differences in results between successively adjusted models reflected the addition of covariates, rather than varying sample size. We also descriptively assessed the degree of overlap between significant factors in each at-risk sample versus the full sample.

### Mendelian randomization (MR) analyses

We performed bidirectional two-sample MR analyses (**Methods S6**) between depression and modifiable factors identified in the fully adjusted factors-wide association scan (Model 2) for the full sample and at-risk groups, if any. For genetic instruments, we accessed the GWAS Atlas online database^24^ (https://atlas.ctglab.nl) to obtain publicly available UK Biobank-based summary statistics for each identified factor. For depression, we retained the summary statistics from the Psychiatric Genomics Consortium^11^ used for polygenic scoring. As genetic instruments, we extracted highly associated SNPs (p < 1×10E-7) that were clumped for independence at r^2^ > 0.001. Using the *TwoSampleMR* package in R^25^, we conducted MR analyses to estimate the effect of each modifiable factor on the risk of depression, and vice versa. For primary MR analyses, we combined per-SNP effects using inverse variance weighted (IVW) meta-analysis, where the resulting estimate represents the slope of a weighted regression of SNP-outcome effects on SNP-exposure effects where the intercept is constrained to zero. We applied MR-PRESSO^26^ in combination with additional tests (i.e., Cook’s distance, studentized residuals, Q-value outliers) to detect statistical outliers reflecting likely pleiotropic bias^27^, and removed these outliers to generate estimates for reporting. We relaxed the instrument p-value threshold for several traits (p < 1×10E-6; i.e., vitamin B; walking frequency) that did not have sufficient SNPs (three or fewer) following outlier removal. We then compared the pattern of IVW results to other established MR methods whose estimates rely on different assumptions and are known to be relatively robust to horizontal pleiotropy, specifically the weighted median approach^28^ and MR Egger regression^29^. We further assessed horizontal pleiotropy using standard methods including leave-one-SNP-out analyses, the modified Cochran’s Q statistic, and the MR Egger intercept test^30^. Finally, for significant results, we searched each instrument SNP in the PhenoScanner v2 database (http://www.phenoscanner.medschl.cam.ac.uk), to identify known associations with depression-related traits at p<1×10E-5 with each instrument SNP or any SNPs that were in linkage disequilibrium at r^2^>0.80, and assessed whether removing these SNPs substantively changed the pattern of results. Reported estimates were converted to odds ratios where the outcome was binary, and interpreted using a conservative Bonferroni-corrected p-value threshold (0.05/number of factors with available summary statistics).

## Results

### Sample description

In the full sample, participants were 54% female and had a mean baseline age of 56.1 (standard deviation, SD = 7.7). 46% reported college or university qualifications, 65% reported current paid or self-employment, and 64% reported average household income of 31,000 pounds or higher. Participants had a mean BMI of 26.7 (SD = 4.4) and 26% endorsed physical illness/disability. Overall, 3.9% met the cut-off for clinically significant symptoms at follow-up; as expected, the prevalence of follow-up depression was elevated within the high PRS group (6.1%) and the high TLE group (12.1%).

### Modifiable factors prospectively associated with depression status in the full sample

In the full sample, 49 factors were significantly associated with depression (Model 0) (**Figure S1 and Table S2a**), ranging across physical activity, media use, sleep, social, environmental, and dietary domains. After adjusting for sociodemographic factors (Model 1), 39 factors were significantly associated with depression (**Figure S2 and Table S2b**). After further adjusting for physical health factors (Model 2), 29 factors remained significantly associated with depression (Figure 2 and **Table S2c**).

**Figure 2.**
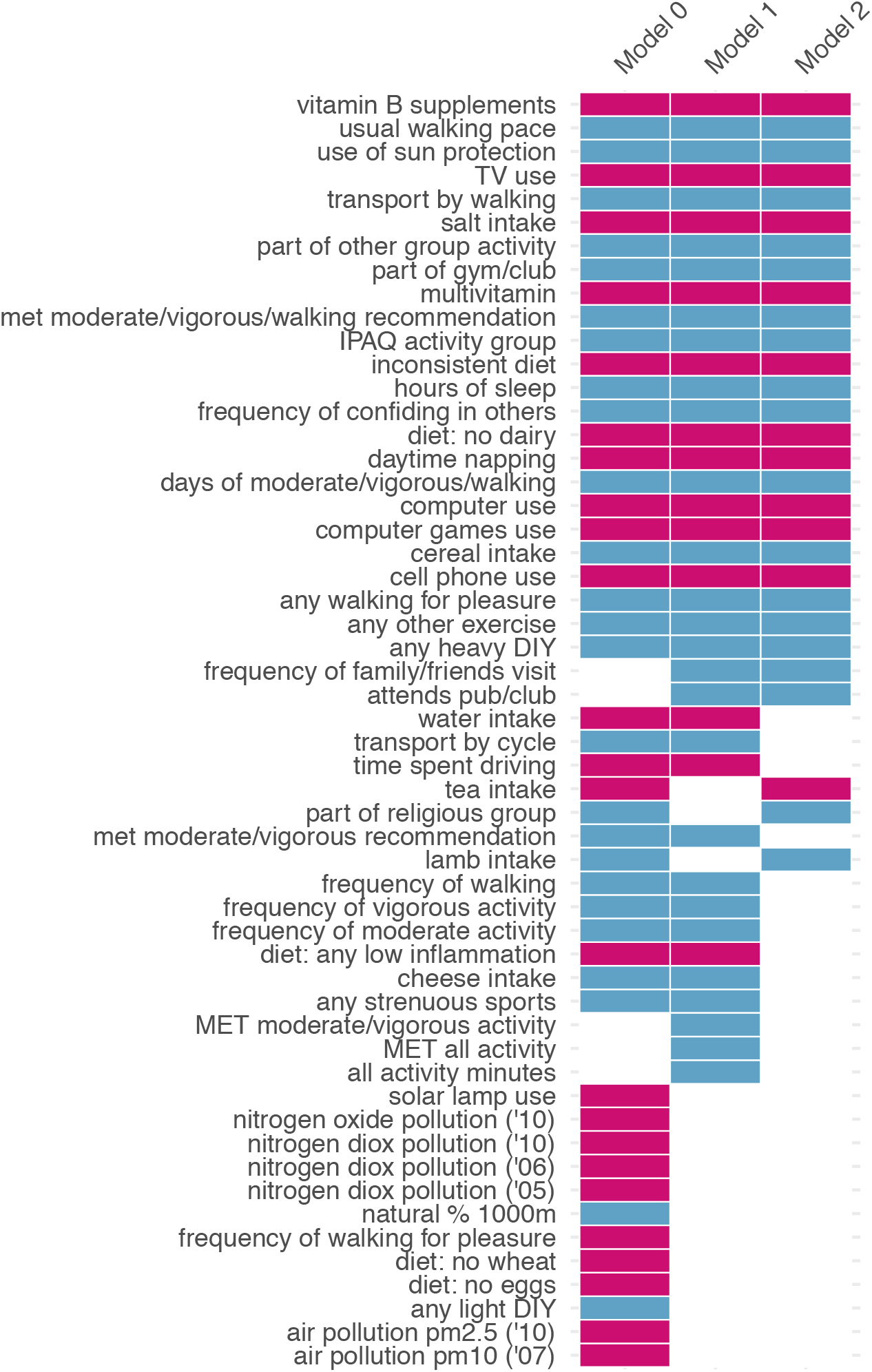
Consistency of top associated factors across levels of covariate adjustment. Blue = reduced odds of depression; red = increased odds of depression. Results are shown only for factors that had significant associations in at least one model.

Of the 29 top hits identified in Model 2, 18 factors were associated with reduced odds of depression and 11 were associated with increased odds of depression (Figure 3; **Table S2c**). The top ten factors included six protective factors: confiding in others (aOR=0.83, 95% CI [0.82-0.85], p=9.50E-100); sleep duration (aOR=0.83 [0.80-0.85], p=5.31E-33); engaging in exercises like swimming or cycling (aOR=0.70 [0.66-0.75], p=2.88E-25); walking pace (aOR=0.79 [0.74-0.84], p=3.33E-15); being part of gym/club (aOR= 0.77 [0.72-0.83]; p=3.93E-12); and cereal intake (aOR=0.89 [0.87-0.92], p=9.58E-12); and four risk factors: daytime napping (aOR=1.29 [1.22-1.37], p=1.20E-19); computer use (aOR=1.10 [1.07-1.13], p=9.38E-12); TV use (aOR=1.12 [1.08-1.16], p=6.05E-12); and cell phone use (aOR=1.10 [1.07-1.13], p=1.25E-11).

**Figure 3.**
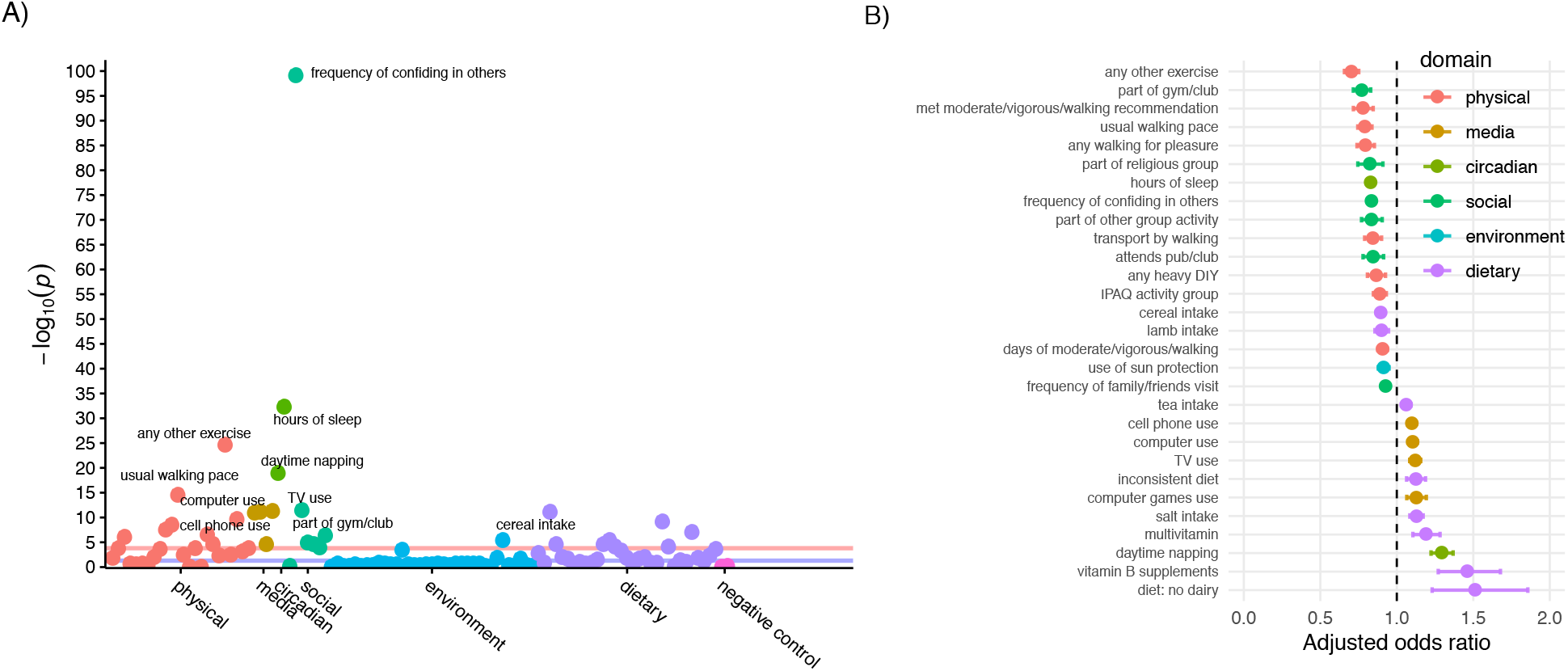
Association results between modifiable factors and clinically significant depression in the full sample, adjusted for sociodemographic and health factors. **A)** Association plot for modifiable factors in relation to follow-up depression, with x-axis organized by conceptual domains, y-axis showing statistical significance as −log10 of p-value, and red horizontal line showing the significance threshold corrected for multiple testing. **B)** Adjusted odds ratios for significant factors, in ascending order (i.e., from risk-reducing to risk-increasing).

### Factors associated with depression among at-risk individuals based on polygenic risk

Among individuals at high predicted probability for depression based on polygenic scores, 12 factors were identified to be significantly associated with depression (Model 0) (**Figure S6 and Table S2d**), ranging across physical activity, media, sleep, social, and dietary domains. After adjustment for sociodemographic factors (Model 1), 10 factors showed associations with depression (**Figure S7 and Table S2e**). After further accounting for physical health factors (Model 2), five factors remained associated with depression (**Figure S9 and Table S2f**). Notably, each of these factors had been identified in the full sample (Figure 2). Of these (**Figure S8**), two were associated with reduced odds of depression: frequency of confiding in others (aOR=0.85 [0.81-0.89], p=2.87E-13) and sleep duration (aOR=0.81 [0.75-0.88], p=4.07E-07). The other two factors were associated with increased odds of depression: time spent using the computer (aOR=1.17 [1.09-1.26], p=1.19E-05), salt intake (aOR=1.21 [1.10-1.30], p=1.31E-04), and time spent using the TV (aOR=1.17 [1.08-1.27], p=1.57E-45).

### Factors associated with depression among at-risk individuals based on traumatic life events

Among individuals with high predicted risk for depression based on their reported traumatic life events, 18 factors were significantly associated with depression (Model 0) (**Figure S13 and Table S2g**). After adjustment for sociodemographic factors (Model 1), 16 factors were significantly associated with depression (**Figure S14 and Table S2h**). After further adjusting for health factors (Model 2), five factors remained associated with depression (**Figure S16 and Table S2i**). Again, each of these factors had been identified in the full sample (Figure 2). Of these (**Figure S15**), four factors were associated with reduced odds of depression: frequency of confiding in others (aOR=0.85 [0.82-0.88], p=5.87E-23); engaging in exercises like swimming or cycling (aOR=0.66 [0.58-0.74], p=6.30E-11); sleep duration (aOR=0.84 [0.79-0.89], p=7.93E-09); and typical walking pace (aOR=0.81 [0.73-0.90], p=1.56E-4), while one factor was associated with increased odds of depression: time spent watching TV (aOR=1.15 [1.08-1.22], p=9.15E-06).

### Follow-up Mendelian randomization (MR) analyses

All modifiable factors identified within at-risk groups had been identified in the full sample; thus, we tested any factors identified in the adjusted full sample (Model 2) with available GWAS summary statistics. Bidirectional MR analyses between each factor and depression revealed a number of findings suggesting causal influences (Figure 4 and Figure 5); results across MR methods for each factor are summarized in **Table S3**.

**Figure 4.**
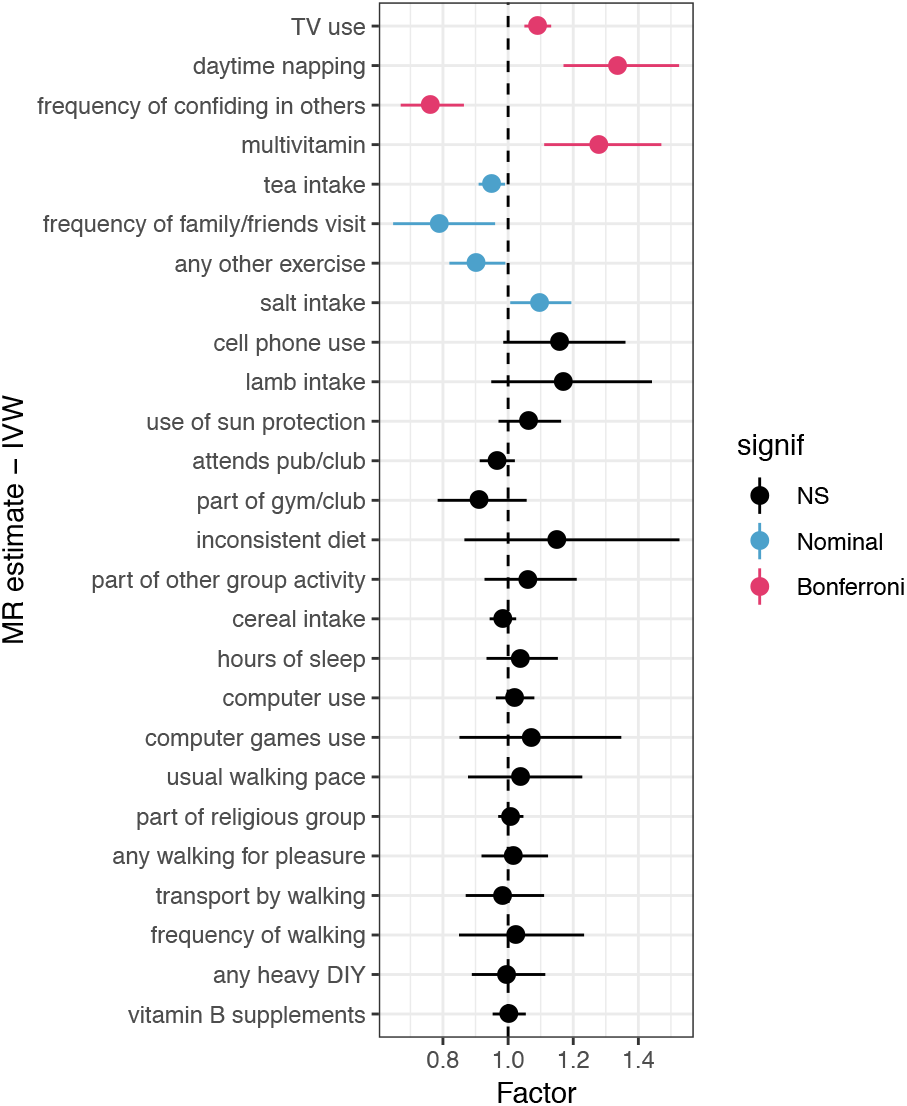
MR estimates of top modifiable factors → the risk of depression with outliers removed, based on the inverse-variance weighted method (for the weighted median method, see Figure S26).

**Figure 5.**
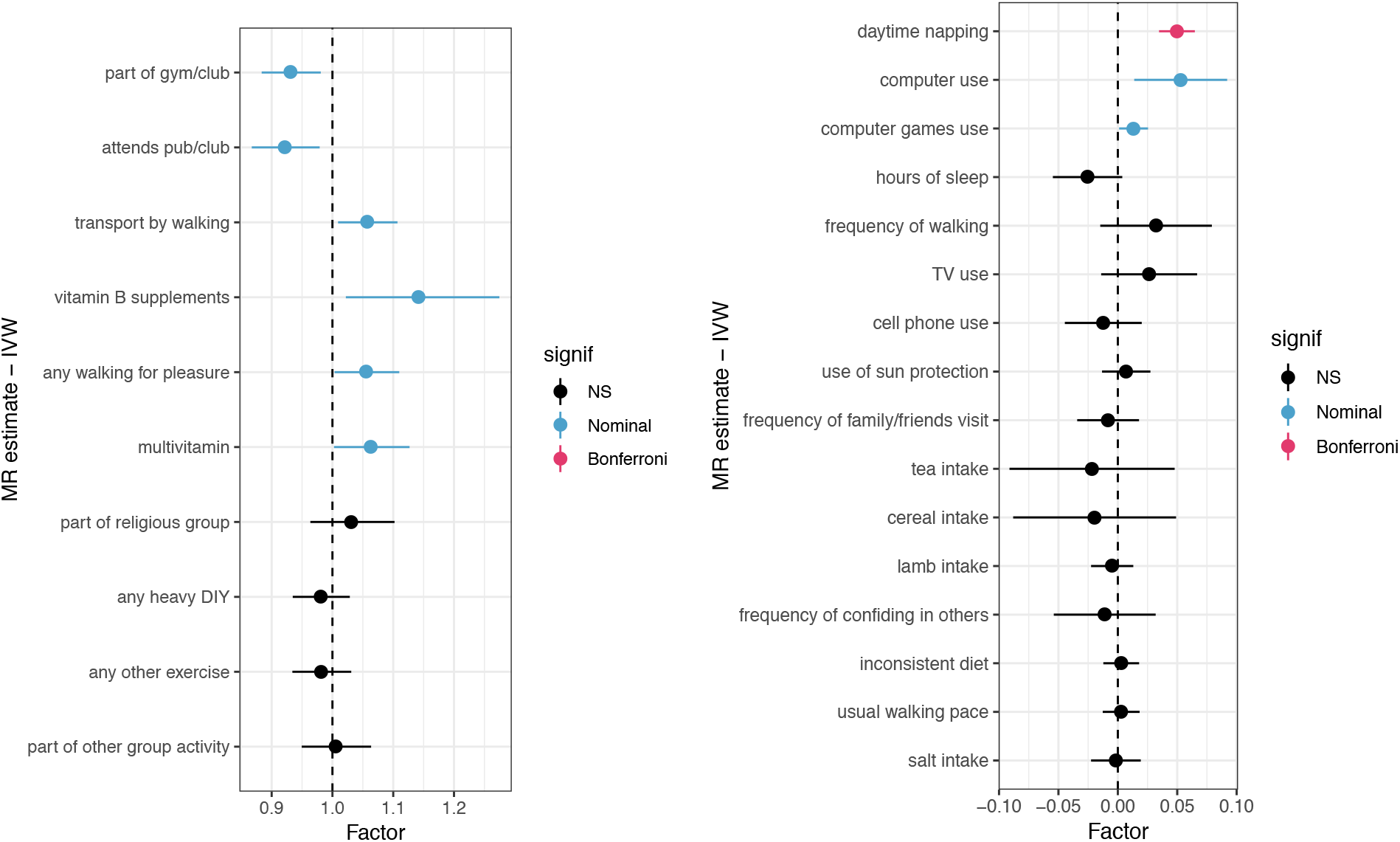
MR estimates of depression → top modifiable factors with outliers removed, based on the inverse-variance weighted method (for the weighted median method, see Figure S27). Odds ratio estimates on left shown for dichotomous factors as outcomes, and beta estimates on right shown for non-dichotomous factors as outcomes.

MR evidence supported a beneficial effect of confiding in others (OR=0.76 [0.67-0.86], p=2.53E-05; 10 SNPs), with non-significant effects in the reverse direction. No effect heterogeneity (Q statistic=5.5, p=0.78) was observed, and the MR-PRESSO global test (p=0.82) and MR Egger intercept test (p=0.78) did not provide evidence of horizontal pleiotropy. Given the lower number of SNPs tested, this effect remained notably significant when relaxing the instrument SNP p-value threshold to 1×10E-6 (OR=0.89 [0.84-0.95], p=7.63E-4; 50 SNPs), and when retaining only SNPs with no known associations with potentially relevant traits in the PhenoScanner database (**Table S3**). We also found MR evidence supporting a deleterious effect of TV use (OR=1.09 [1.04-1.15], p=1.33E-03; 145 SNPs), with non-significant effects in the reverse direction. No effect heterogeneity (Q statistic=130.8, p=0.78) was observed, and the MR-PRESSO global test (p=0.44) and MR Egger intercept test (p=0.78) did not provide evidence of horizontal pleiotropy.

Daytime napping showed bidirectional effects with depression, such that daytime napping was linked to higher odds of depression (OR=1.34 [1.17-1.53], p=1.82E-05; 91 SNPs) while depression was also associated with increased daytime napping (beta=0.05 [0.03-0.06], p=8.45E-11; 43 SNPs), with no evidence of effect heterogeneity or horizontal pleiotropy in either direction. Surprisingly, MR evidence suggested that multivitamin use was also linked to increased odds of depression (OR=1.28 [1.11-1.47], p=6.04E-04; 6 SNPs); however, with the lower number of SNPs tested, this effect was notably attenuated when relaxing the instrument SNP p-value threshold to 1×10E-6 (OR=1.07 [1.0-1.14], p=0.0498; 30 SNPs). Depression was also nominally associated with increased intake of multivitamins (OR=1.06 [1.002-1.13], p=4.07E-2; 44 SNPs). Other nominal results at the traditional p<0.05 threshold are summarized in **Methods S7** and included tea intake (protective), frequency of family/friend visits (protective), exercises such as cycling or swimming (protective), and salt intake (risk-increasing).

## Discussion

Although depression is recognized as the leading cause of disability worldwide, decades of research have identified few actionable prevention strategies. We recently used MR to validate the protective effect of objectively measured physical activity^5^ on depression risk, but have expanded this program of research to assess contributions to depression risk from a wider range of modifiable factors. Using phenotypic and genomic data from UK Biobank participants, we screened a broad panel of potentially modifiable factors that may offset genetic and environmental vulnerability to incident depression. Consistent with literature regarding the multifactorial nature of depression risk^31^, we identified numerous modifiable factors that were prospectively associated with depression ranging across multiple domains, including physical activity, social, media-related, and dietary domains.

Several factors identified to be protectively associated with depression were not unexpected—but received novel validation in an MR framework. The frequency of confiding in others, an index of emotional social support, showed a robust phenotypic association consistent with MR results even after removing potentially pleiotropic instruments, suggesting that increased confiding in others is likely to be causally linked with reduced odds of depression. This echoes previous evidence on the mood-related benefits of social support^6^ and substantiates its role in preventing depression. Frequency of visiting family/friends also showed nominal results in the MR framework, suggesting that multiple dimensions of social support (not only reaching out to confide in others, but generally having increased social interactions) may help reduce risk of depression. Consistent with these findings, we previously demonstrated that greater levels of social cohesion in a sample of military personnel reduced the risk of incident depression despite high polygenic risk or exposure to traumatic events^32^.

As expected, engagement in various kinds of physical activity showed protective associations with follow-up depression. However, these relationships were not strongly supported in the MR framework. We previously observed^5^ using MR that while objective measures of physical activity (not included here) are linked to reduced odds of depression, self-report measures of physical activity do not necessarily show these patterns. Objective measures of physical activity—reflecting a broad tendency for movement/activity—have demonstrated higher heritability^33^ and may yield more powerful genetic instruments in an MR framework. Indeed, the self-report activity variables examined in the present study tended to have fewer genome-wide significant SNPs than other traits examined (e.g., media use).

Our MR findings were also consistent with prior evidence that increased TV time is a risk factor for poor mental health, including depression^34^. This work adds to broader evidence linking screen time to mental health outcomes^35^ and suggests that reducing hours spent watching TV could have a role in mitigating depression risk. It remains unclear whether these effects are due to screen time and media exposure per se, or whether TV watching serves as a proxy for time spent in sedentary behavior more generally, which has been associated with depression^36^. Nominal MR results also suggested some possibility for reverse causation, where increased use of other screen-based media (e.g., computer use) appeared to be a consequence of depression.

Other modifiable factors that have received less attention also emerged in our scan. Daytime napping was not only phenotypically associated with increased depression risk but also showed bidirectional influences in the MR context. These results suggest that increased daytime napping could increase risk of depression, and therefore limiting (excessive) daytime napping could prove beneficial for reducing depression risk—however, that depression may also increase one’s tendency to engage in daytime napping. Conversely, greater overall sleep time was associated with reduced odds of follow-up depression; however, this relationship was not substantiated in the MR framework. This may be because sleep shows a more complex and non-linear relationship with depression^37^ than could be modeled in this broad focus study.

Among the more surprising findings was an association between multivitamin use and increased odds of follow-up depression that was supported by initial MR analysis, though attenuated when considering a more relaxed set of genetic instruments. We also found evidence of reverse causation (whereby individuals with depression may turn to vitamin supplementation), which may have contributed to this phenotypic association. To our knowledge, no studies have linked multivitamin use to increased risk of depression so caution is required, particularly since a meta-analysis of multivitamin supplementation trials found no significant effect on depressive symptoms^38^. Prior studies of vitamin intake and depression have largely focused on individual micronutrients (such as vitamin D and folate), with highly mixed results^39^.

Environmental factors such as pollution and exposure to natural environment showed initial associations with depression—risk and protective effects, respectively—that did not persist after adjusting for sociodemographic factors, and were thus not tested in the MR framework. While these unadjusted findings were in line with growing evidence^40^, it may be that environmental exposures exert stronger influences earlier in development^41^, or shape lifetime mental health risk rather than incident cases in a relatively short follow-up period. In addition, sub-features of the natural environment (e.g., tree versus grass coverage) have shown divergent effects on mental health risk^42^, requiring more nuanced study than possible here.

The recent emergence of precision medicine as a framework for disease prevention and treatment motivated our effort to examine whether protective factors operate differentially among individuals at higher risk for depression. We found that all factors identified as protective among at-risk individuals (whether based on polygenic risk or reported traumatic life events) were also protective in the full sample. This suggests that the benefits of these identified factors can be observed in the presence of static vulnerability factors such as genetic risk or reported traumatic life events.

Our study should be evaluated in light of several limitations. First, while we considered a wide array of lifestyle and environmental factors, we were limited by the variables available in the UK Biobank database, which did not include coping styles or psychological factors that could also be modifiable with respect to depression risk. Second, our study relied on self-report measures which can be subject to reporting biases and may not optimally capture all underlying factors of interest. Our assessment of follow-up depression was based on a self-report measure that, while widely used, may not be fully concordant with a clinical diagnostic interview. In addition, the self-reported outcome could explain stronger associations with factors that were also self-reported and have an emotional component (e.g., social factors). Third, confirmation of causal effects may require randomized controlled trials of preventive interventions. In some cases, such trials might be prohibitively costly, require long duration of follow-up, or be otherwise unfeasible. We sought to address this with a prospective design complemented by Mendelian randomization analyses, which provides an important alternative for verifying actionable strategies. However, it should be noted that MR estimates reflect lifelong average effects of genetic variants and should not be interpreted in the same way as effects from a discrete intervention trial or within a brief follow-up period. The absence of an MR result does not disconfirm the potential importance of a factor operating within more acute time frames. Finally, our analyses were restricted to an older white British sample that volunteered for research and thus represents a healthier and more engaged population^43^, and may thus not be generalizable to other populations.

In conclusion, there has to date been limited systematic, large-scale research on modifiable factors for depression. Here, we demonstrate a novel two-stage approach to identifying factors that may protect against the development of depression. Using a large-scale sample with both genomic and wide-ranging lifestyle and environmental measures, we screened more than 100 potentially modifiable factors for their association with incident depression, including among at-risk individuals, and then tested potential causal effects in a Mendelian randomization framework. Our results prioritize an array of potential targets for prevention strategies, including behavioral and lifestyle factors, aimed at reducing the risk of depression. In light of the enormous burden of depression and the dearth of validated avenues for prevention, our results may have important implications for the future of precision psychiatry.

## Acknowledgements

K.W.C. was supported in part by a NIMH T32 Training Fellowship (T32MH017119). J.W.S is a Tepper Family MGH Research Scholar and supported in part by the Demarest Lloyd, Jr, Foundation. T.G. is supported in part by NIA grant K99AG054573. J.R.I.C and G.B. are funded partly by the UK National Institute of Health Research (NIHR), and partly by a grant from Cohen Veterans Bioscience. This paper represents independent research funded in part by the NIHR Biomedical Research Centre at South London and Maudsley NHS Foundation Trust and King’s College London. The views expressed are those of the authors and not necessarily those of the NIH, NHS, NIHR or the Department of Health and Social Care. This research was conducted using the UK Biobank resource under an approved data request (#32568). This work involved the use of the Enterprise Research Infrastructure & Services (ERIS) at Partners HealthCare.

We would like to thank the research participants and employees of 23andMe, Inc. for making this work possible. The following members of the 23andMe Research Team contributed to this study: Michelle Agee, Babak Alipanahi, Adam Auton, Robert K. Bell, Katarzyna Bryc, Sarah L. Elson, Pierre Fontanillas, Nicholas A. Furlotte, Barry Hicks, David A. Hinds, Karen E. Huber, Ethan M. Jewett, Yunxuan Jiang, Aaron Kleinman, Keng-Han Lin, Nadia K. Litterman, Jennifer C. McCreight, Matthew H. McIntyre, Kimberly F. McManus, Joanna L. Mountain, Elizabeth S. Noblin, Carrie A.M. Northover, Steven J. Pitts, G. David Poznik, J. Fah Sathirapongsasuti, Janie F. Shelton, Suyash Shringarpure, Chao Tian, Joyce Y. Tung, Vladimir Vacic, Xin Wang, Catherine H. Wilson

