## Supplementary Figures for "A two-stage approach to identifying and validating modifiable factors for the prevention of depression"

### Modifiable factors associated with depression in the full sample

### Figure S1. Top hits for Model 0 (adjusted for base factors)

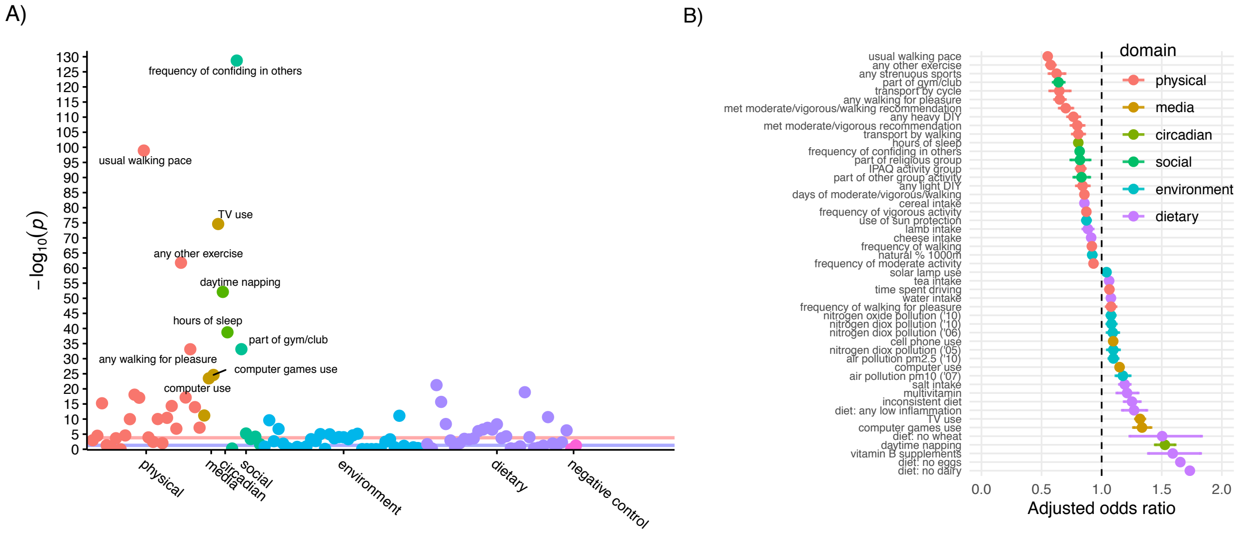


### Figure S2. Top hits for Model 1 (adjusted for sociodemographic factors)

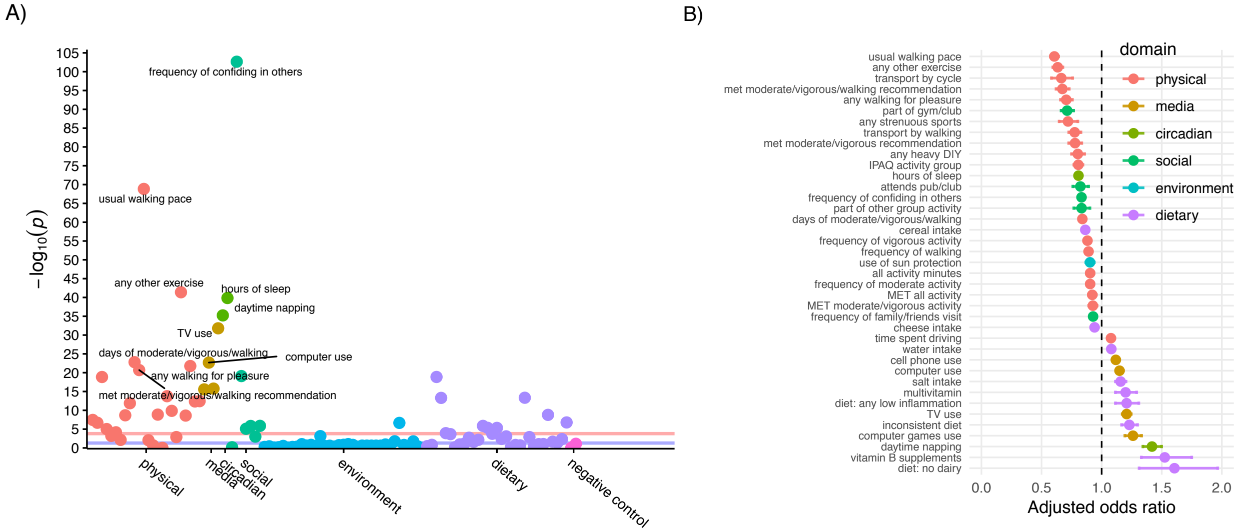


### Top hits for Model 2 (further adjusted for sociodemographic and health factors)

*Shown in main manuscript*

**Figure S3. All results for Model 0 (adjusted for base factors)**

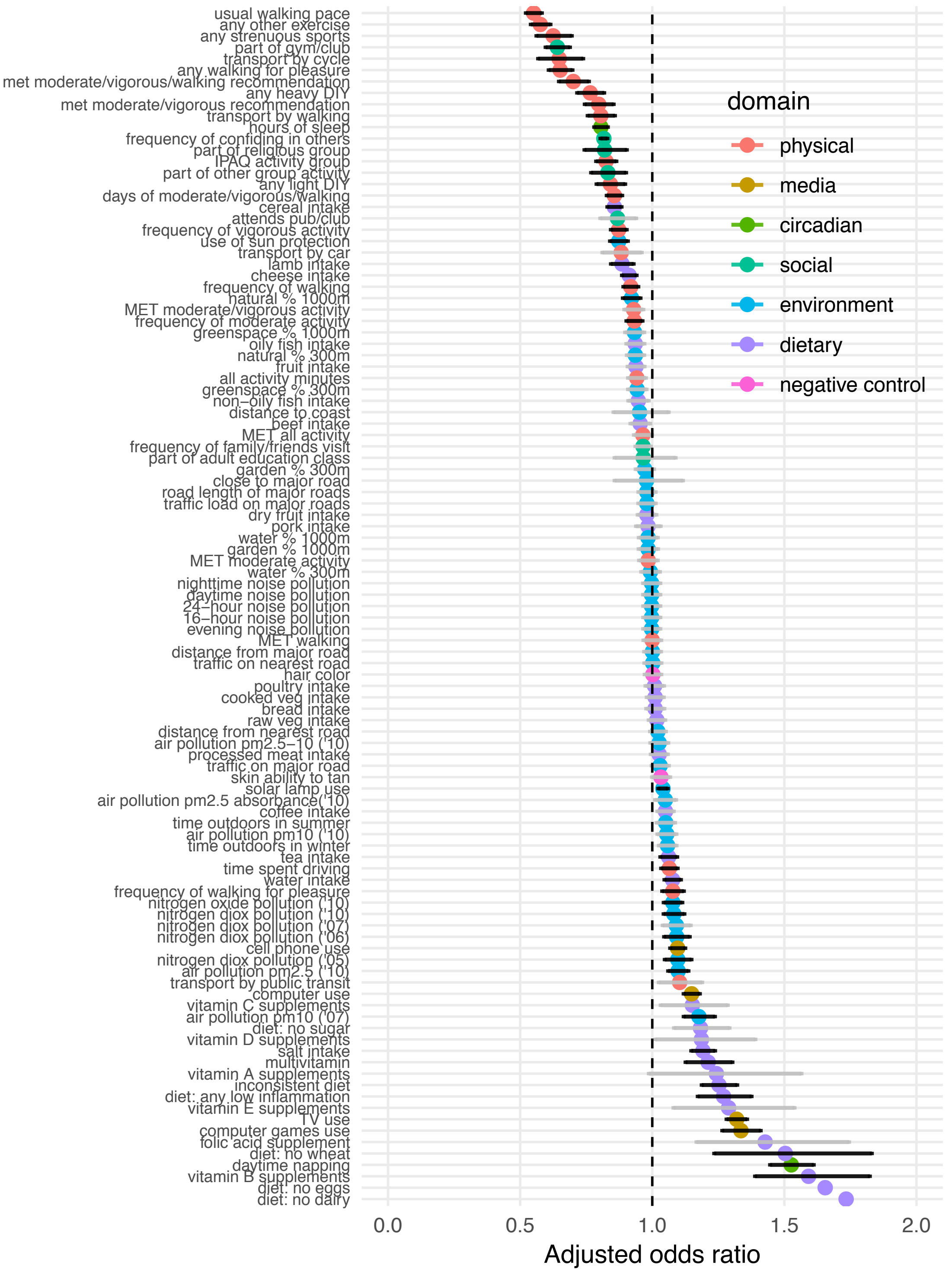

**Figure S4. All results for Model 1 (further adjusted for sociodemographic factors)**

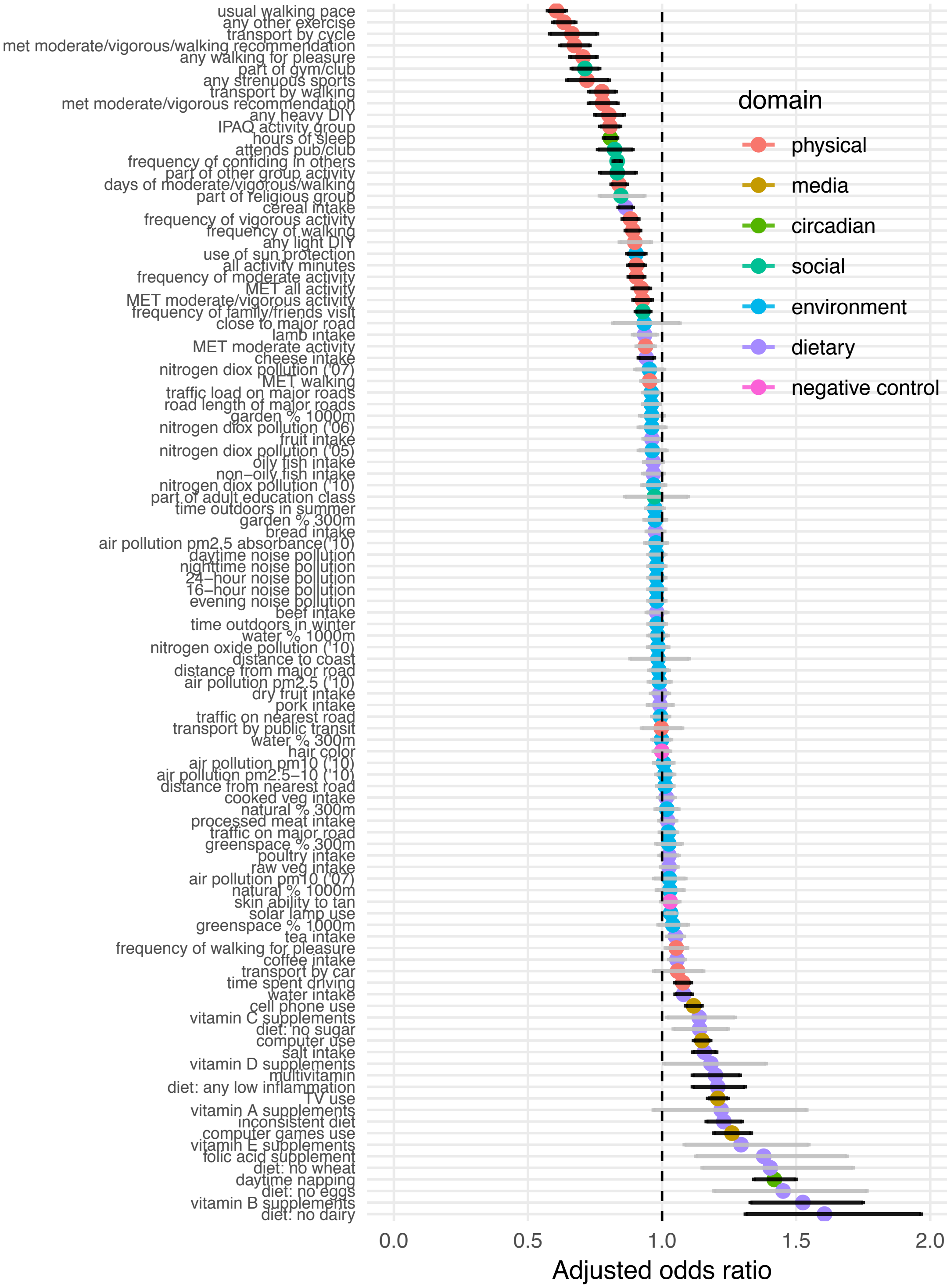

**Figure S5. All results for Model 2 (further adjusted for sociodemographic and health factors)**

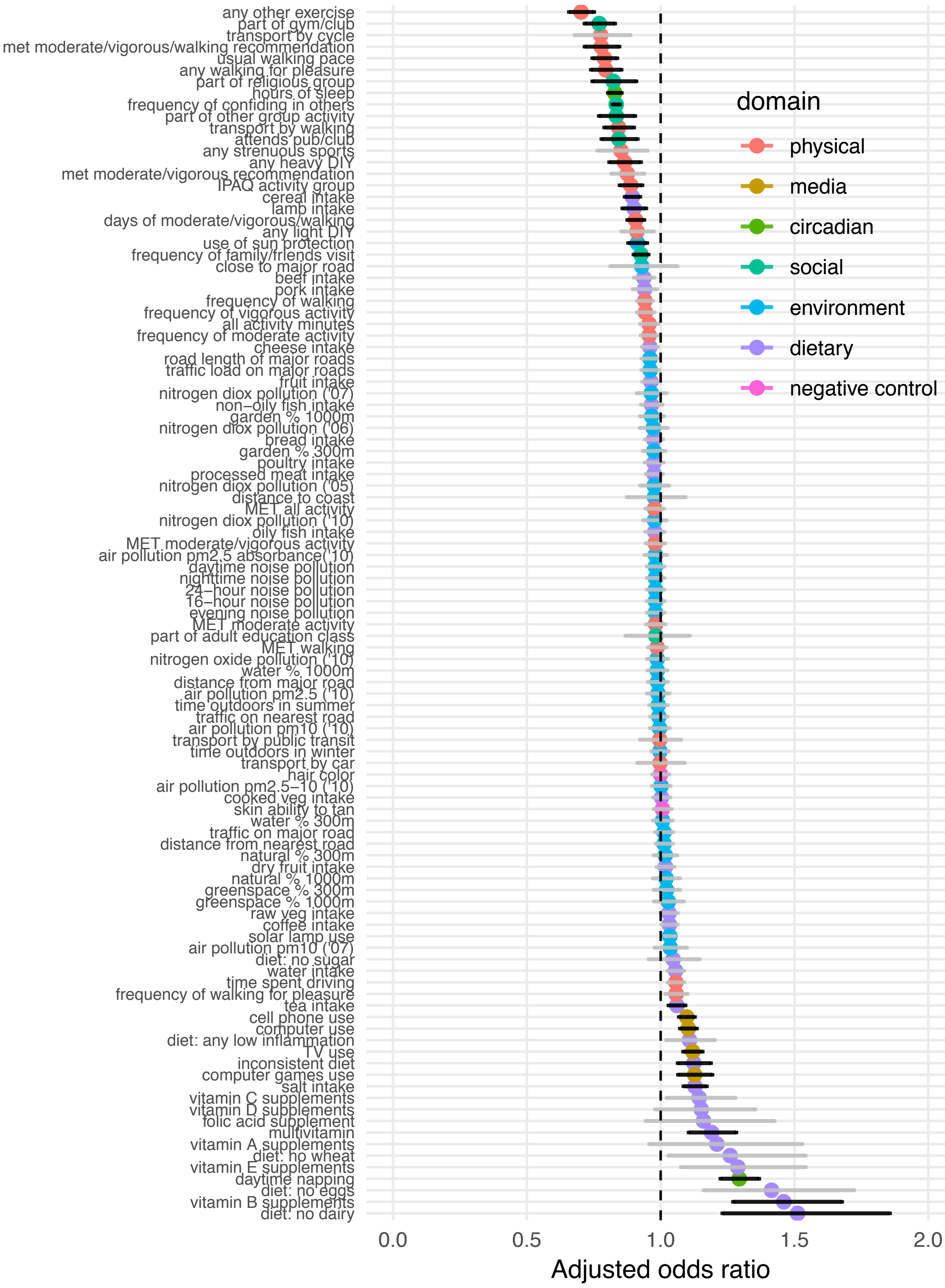

### Factors associated with depression among at-risk individuals (based on polygenic risk)

#### Figure S6. Top hits for Model 0 (adjusted for base factors)

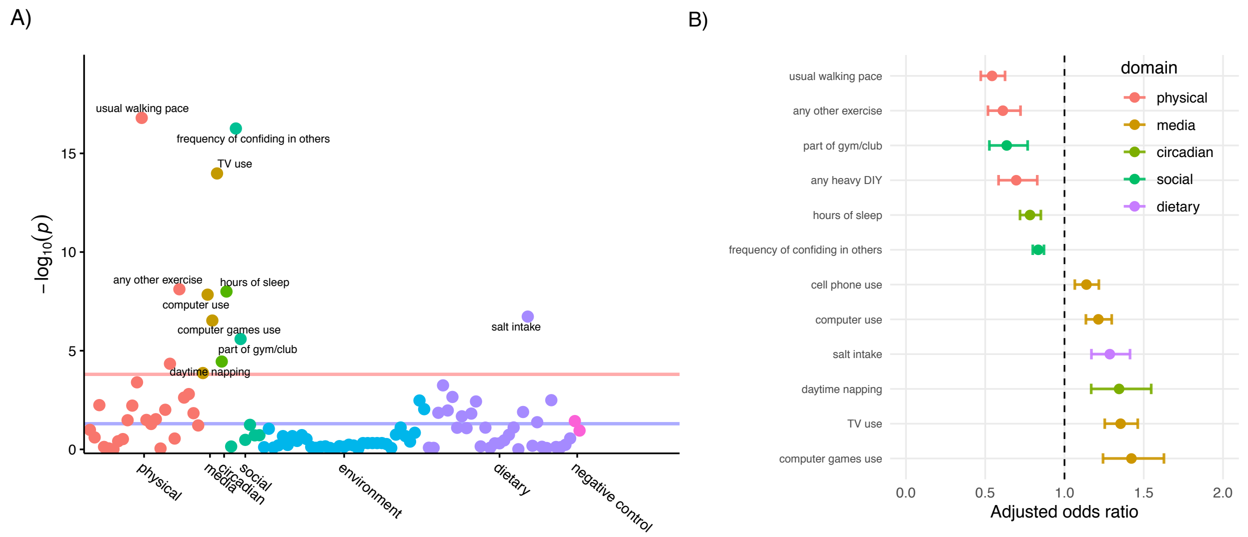


#### Figure S7. Top hits for Model 1 (further adjusted for sociodemographic factors)

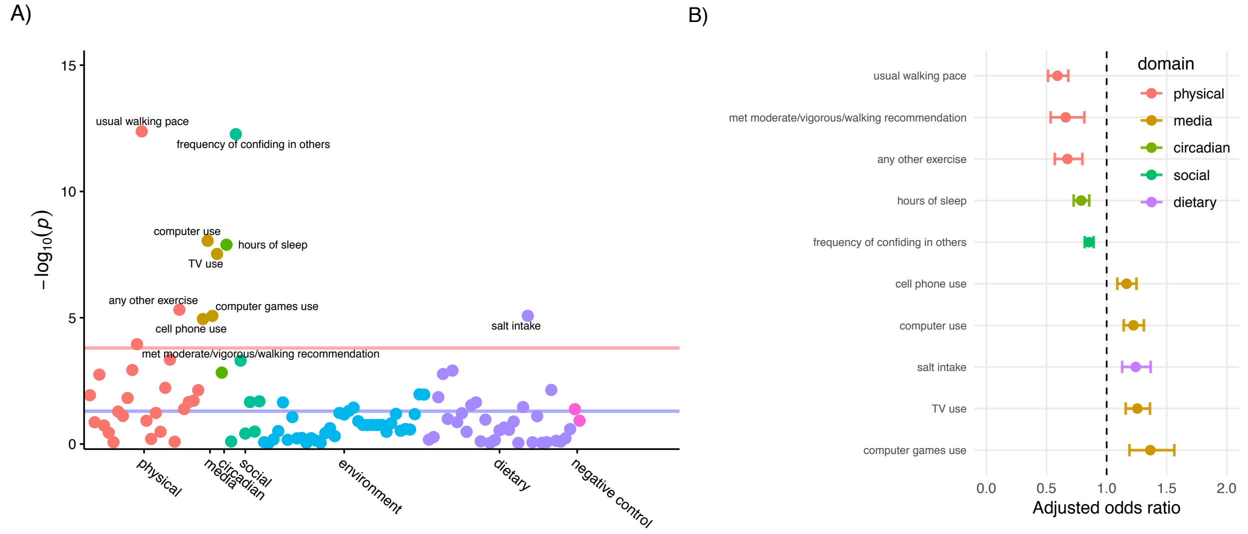

**Figure S8. Top hits for Model 2 (further adjusted for sociodemographic and health factors)**

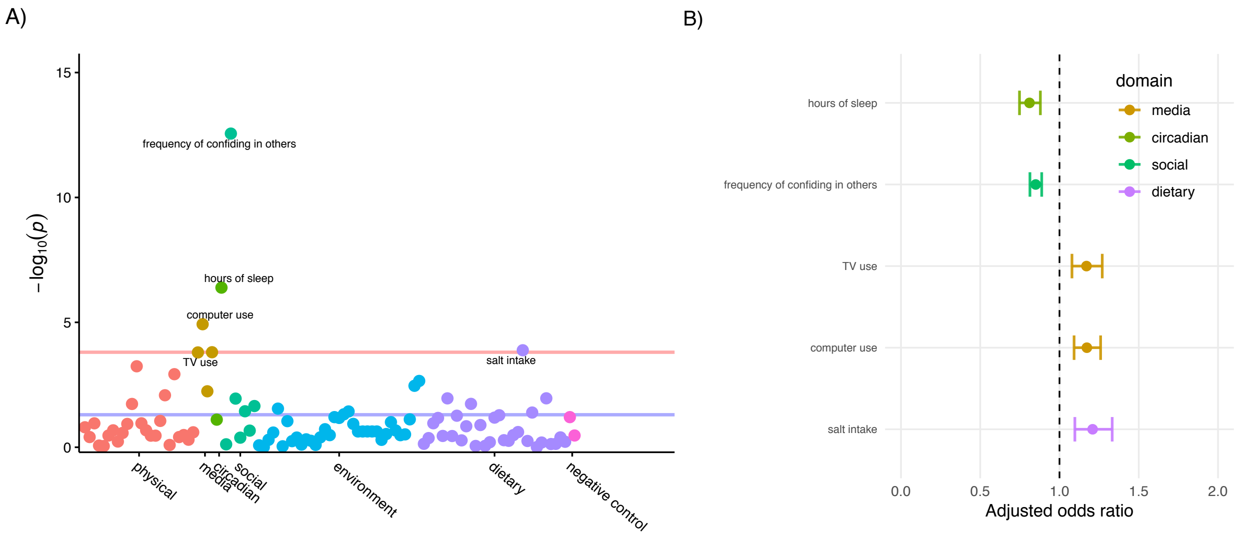


**Figure S9. Consistency of associated factors across levels of covariate adjustment.** Blue = protective direction of association; red = risk-increasing direction of association.

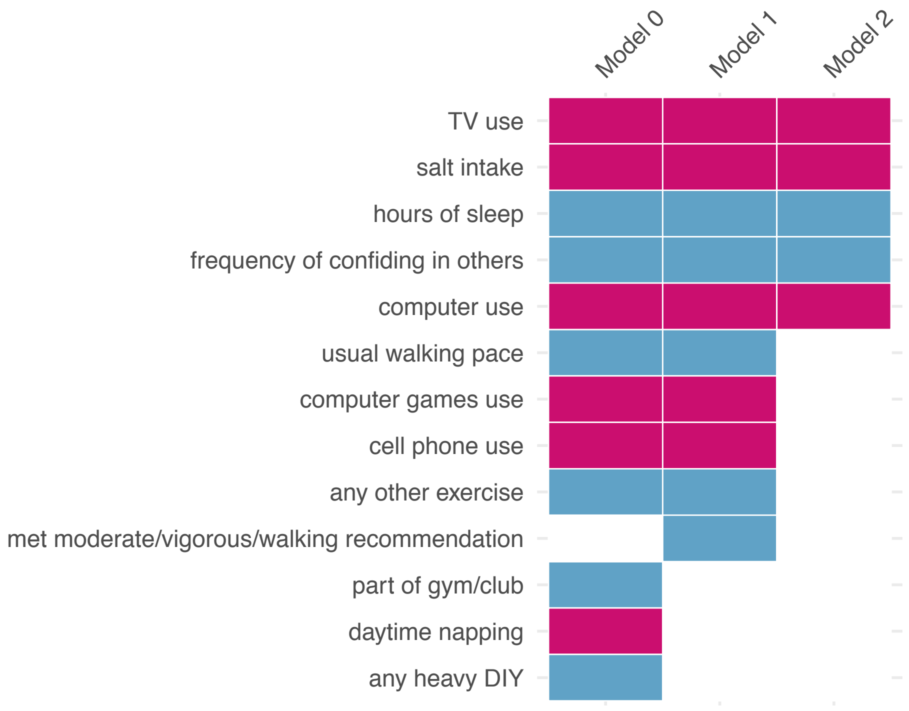

**Figure S10. All results for Model 0 (adjusted for base factors)**

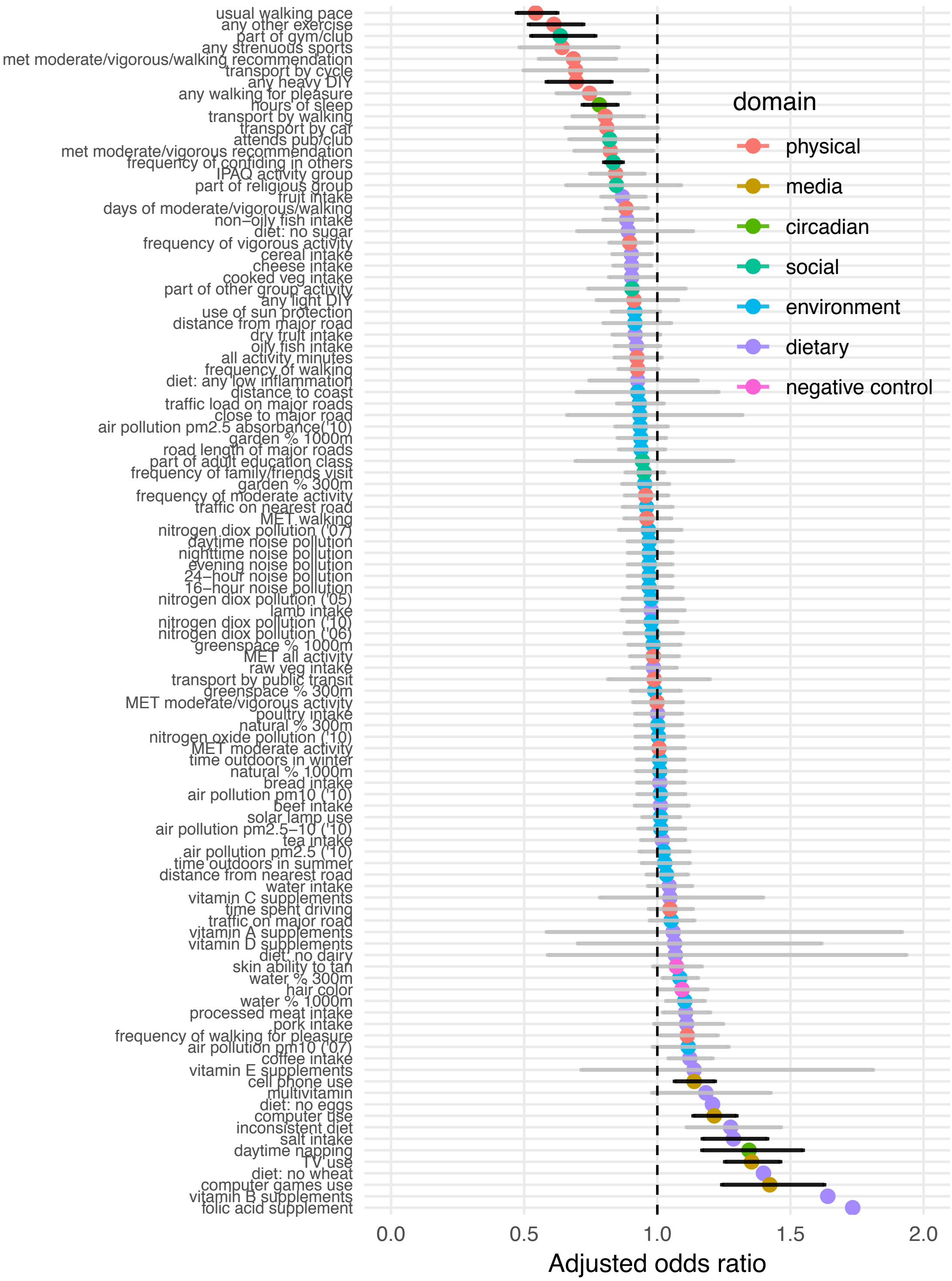

**Figure S11. All results for Model 1 (further adjusted for sociodemographic factors)**

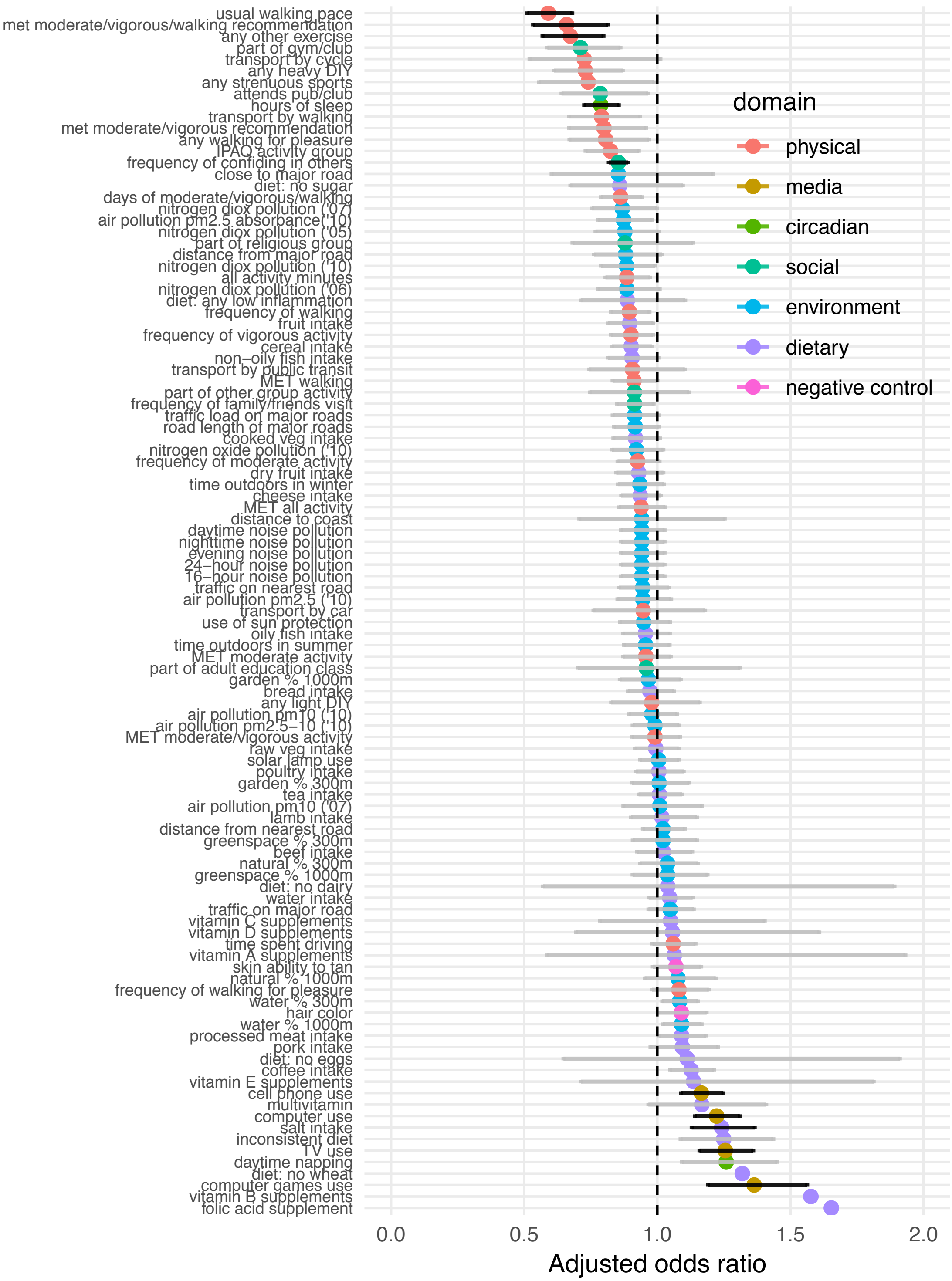

**Figure S12. All results for Model 2 (further adjusted for sociodemographic and health factors)**

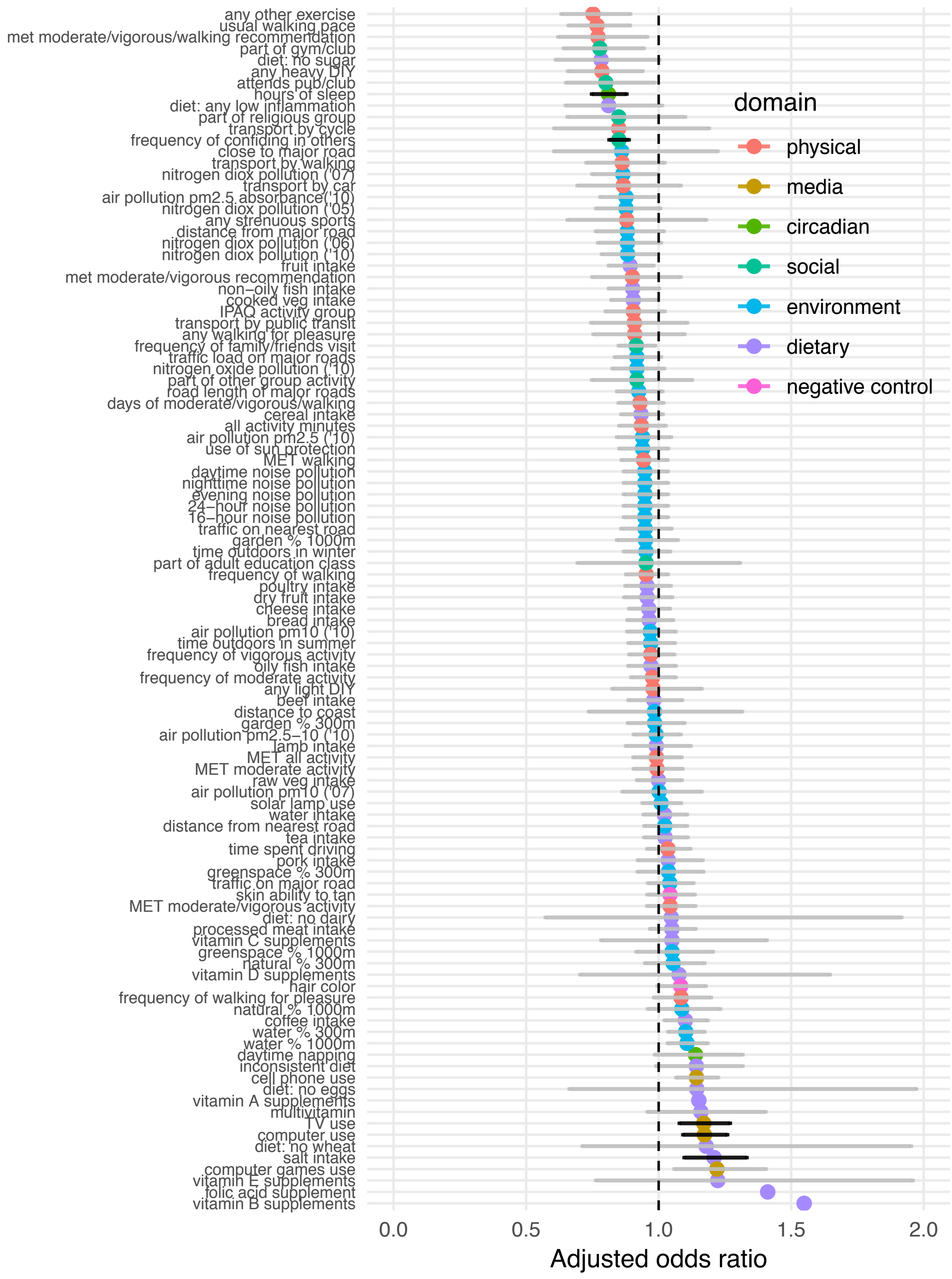

**Factors associated with depression among at-risk individuals (based on traumatic life events)**

**Figure S13. Top hits for Model 0 (adjusted for base factors)**

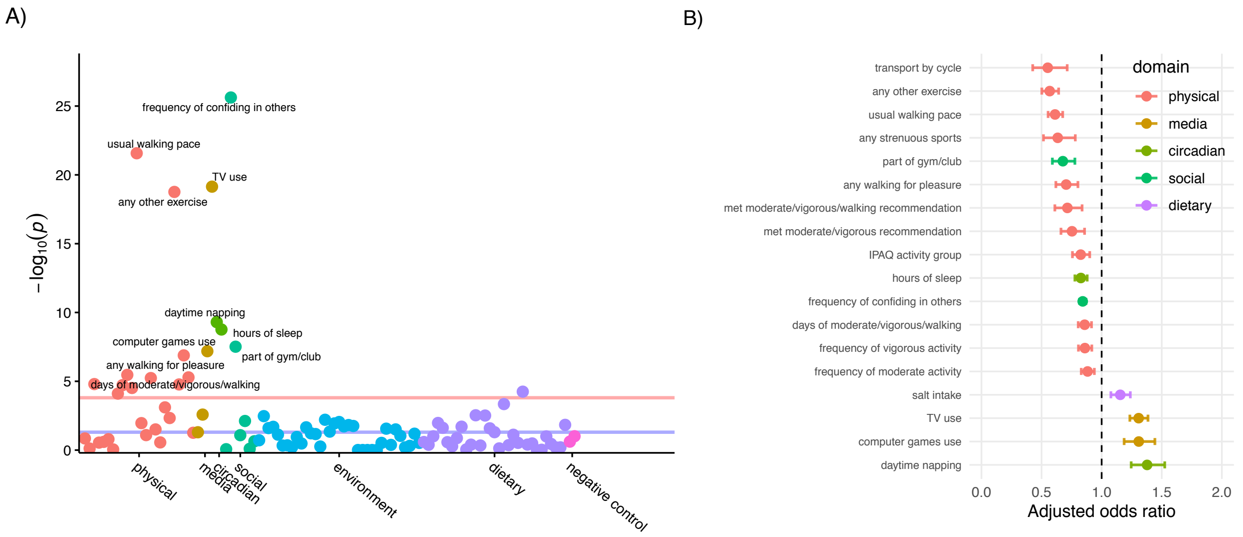


**Figure S14. Top hits for Model 1 (further adjusted for sociodemographic factors)**

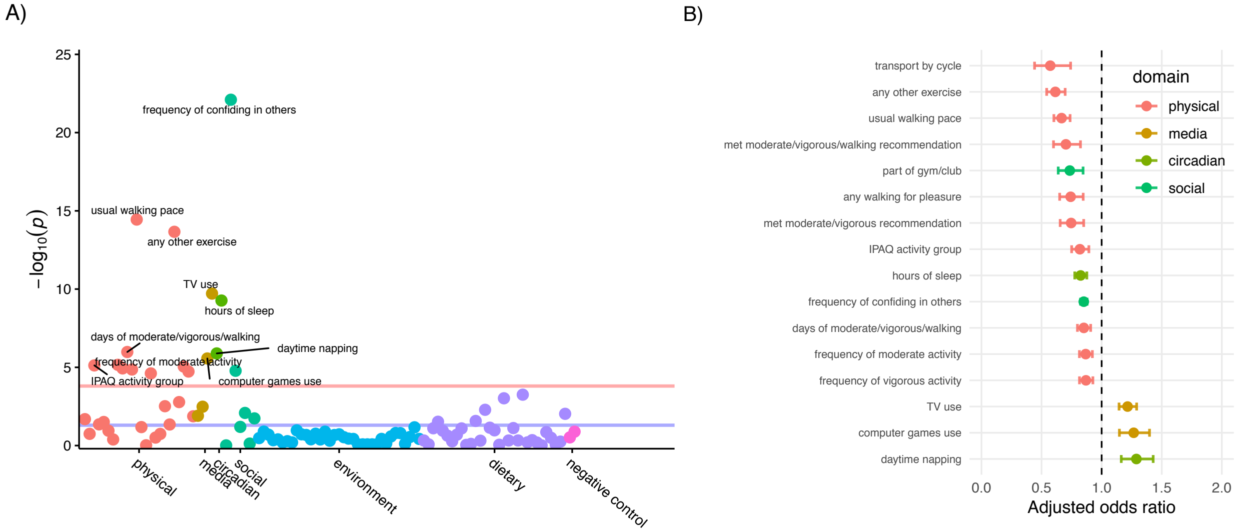

**Figure S15. Top hits for Model 2 (further adjusted for sociodemographic and health factors)**

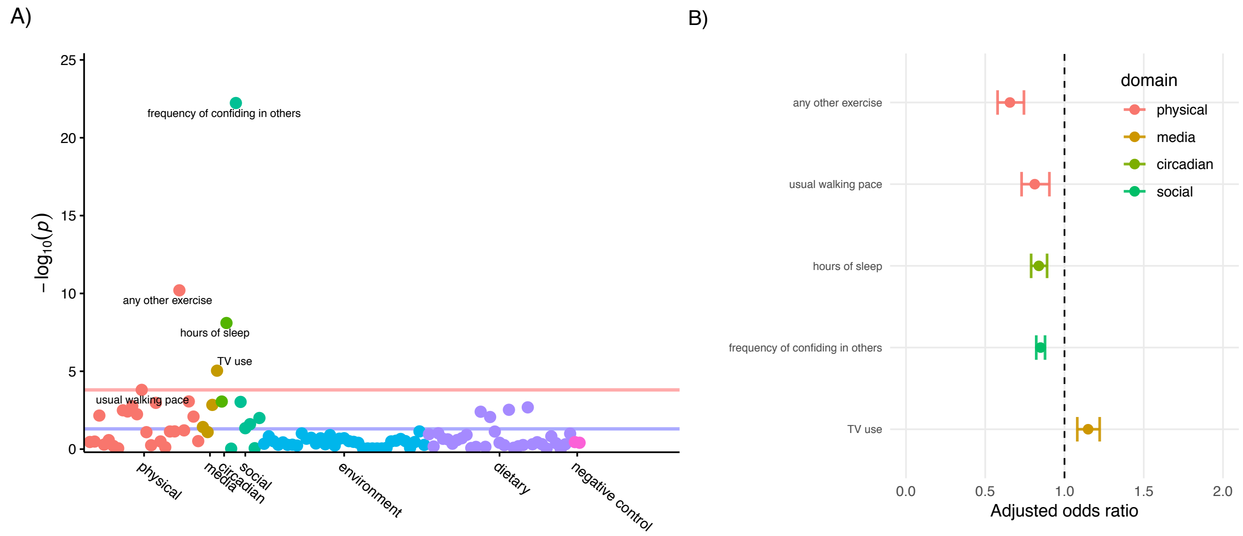


**Figure S16. Consistency of associated factors across levels of covariate adjustment.** Blue = protective direction of association; red = risk-increasing direction of association.

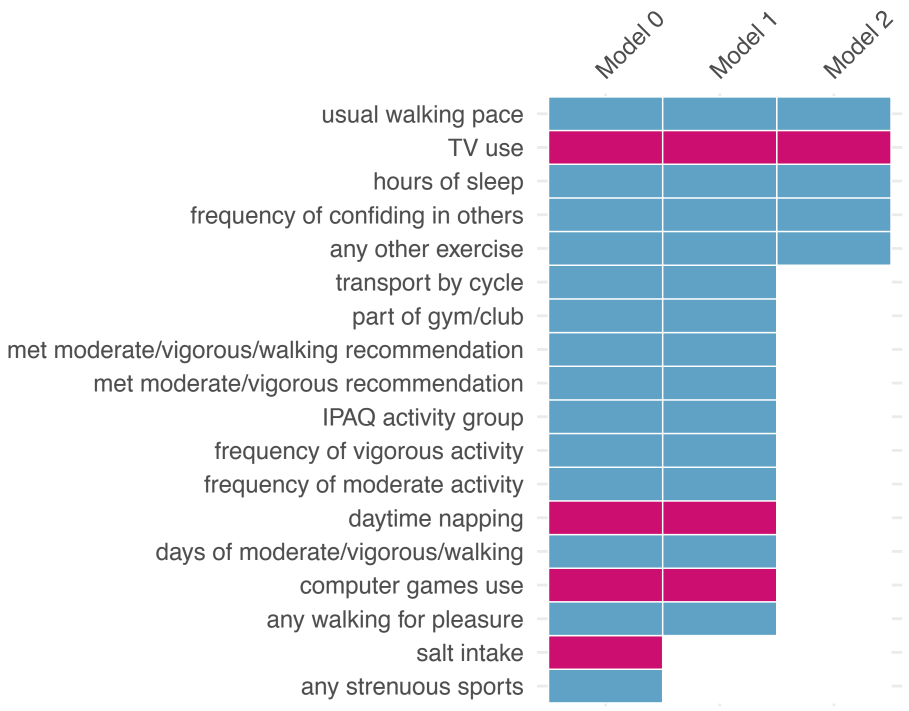

**Figure S17. All results for Model 0 (adjusted for base factors)**

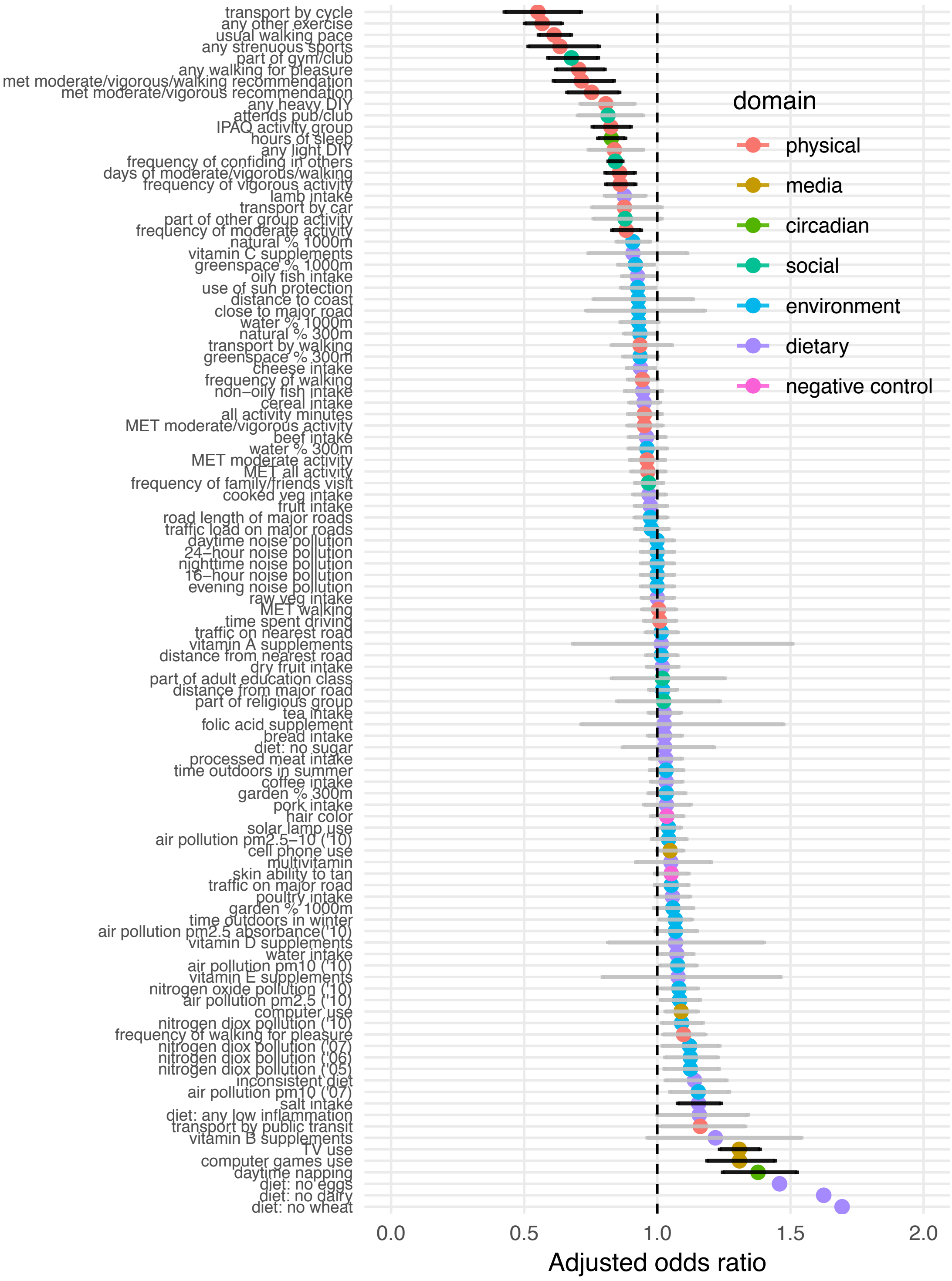

**Figure S18. All results for Model 1 (further adjusted for sociodemographic factors)**

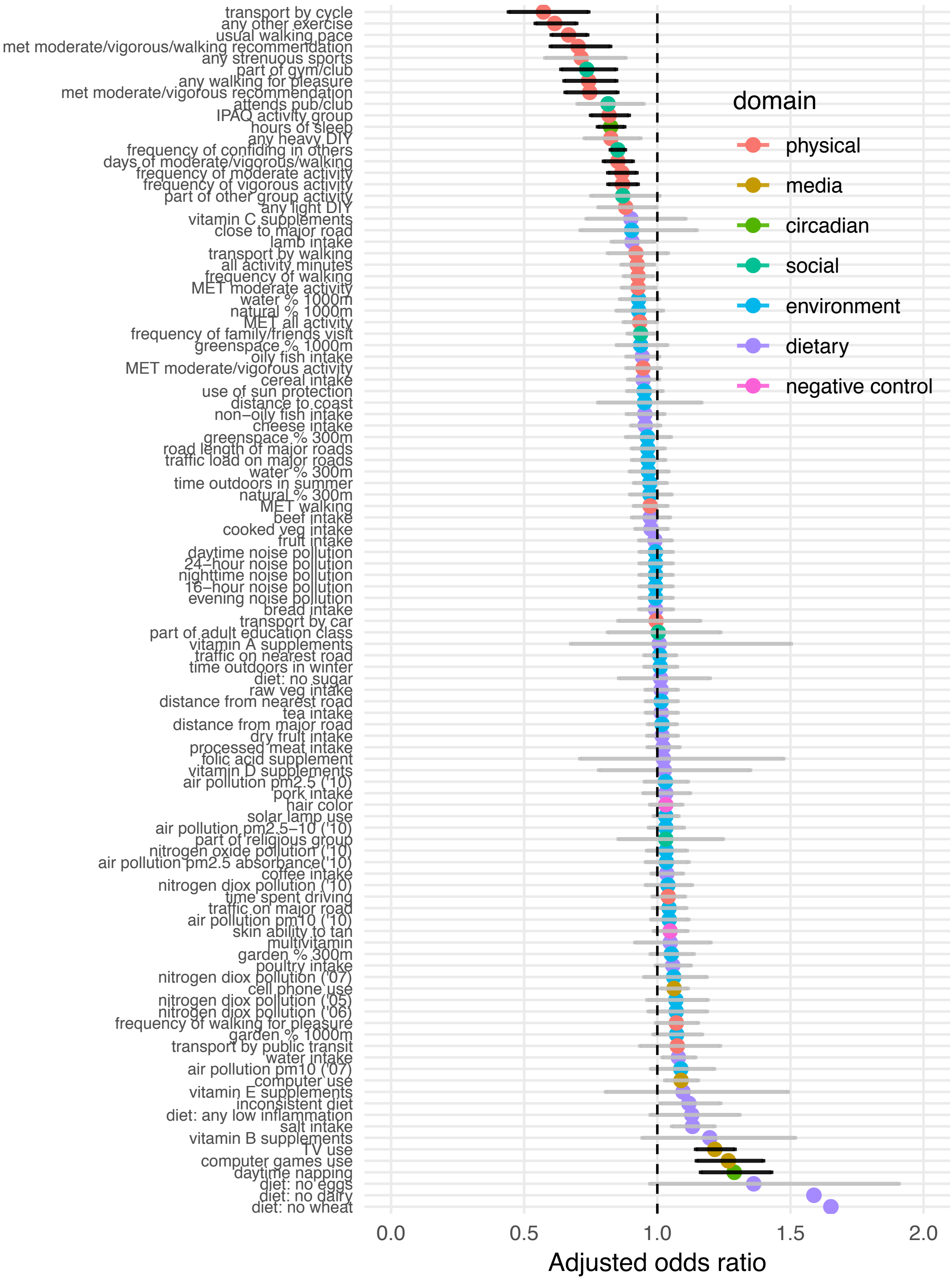

**Figure S19. All results for Model 2 (further adjusted for sociodemographic and health factors)**

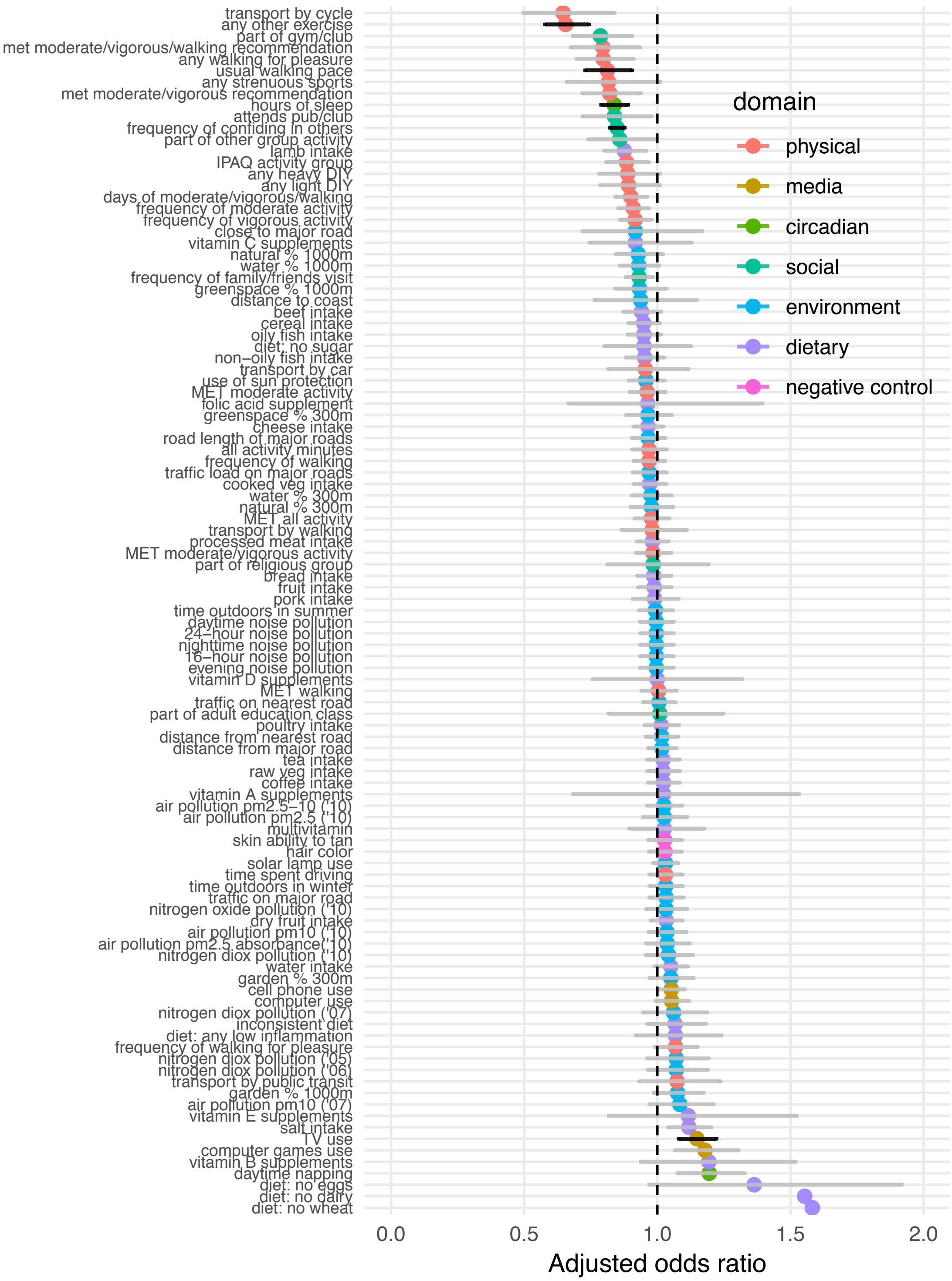

**Exploratory factors associated with depression among at-risk individuals (based on being at-risk on both polygenic risk and traumatic life events, maximum n=1,558)**

**Figure S20. Top hits for Model 0 (adjusted for base factors)**

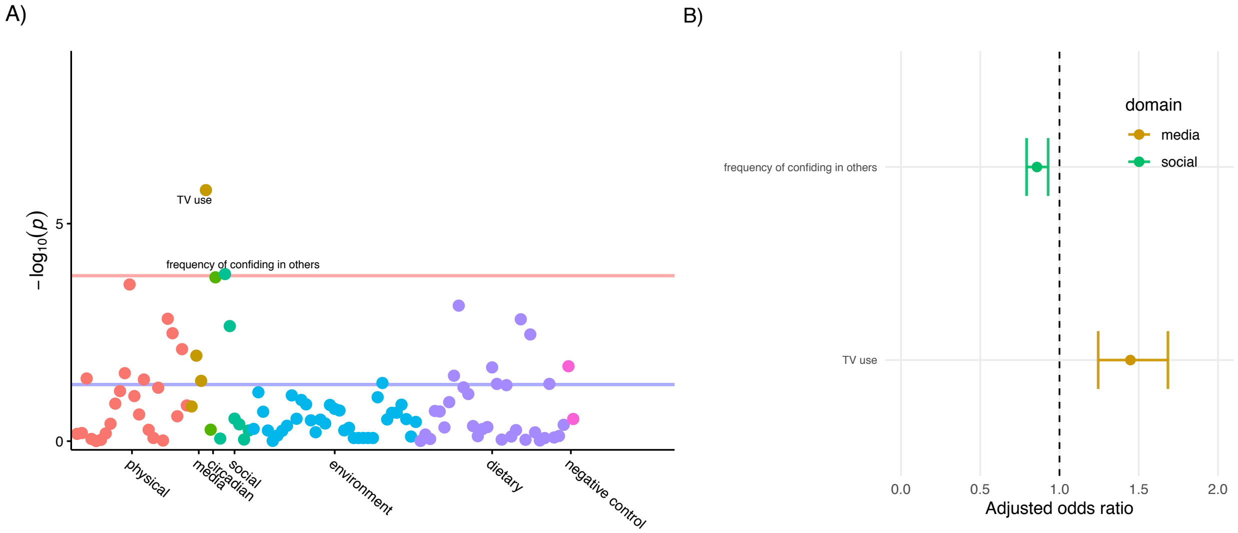


**Figure S21. Top hits for Model 1 (further adjusted for sociodemographic factors)**

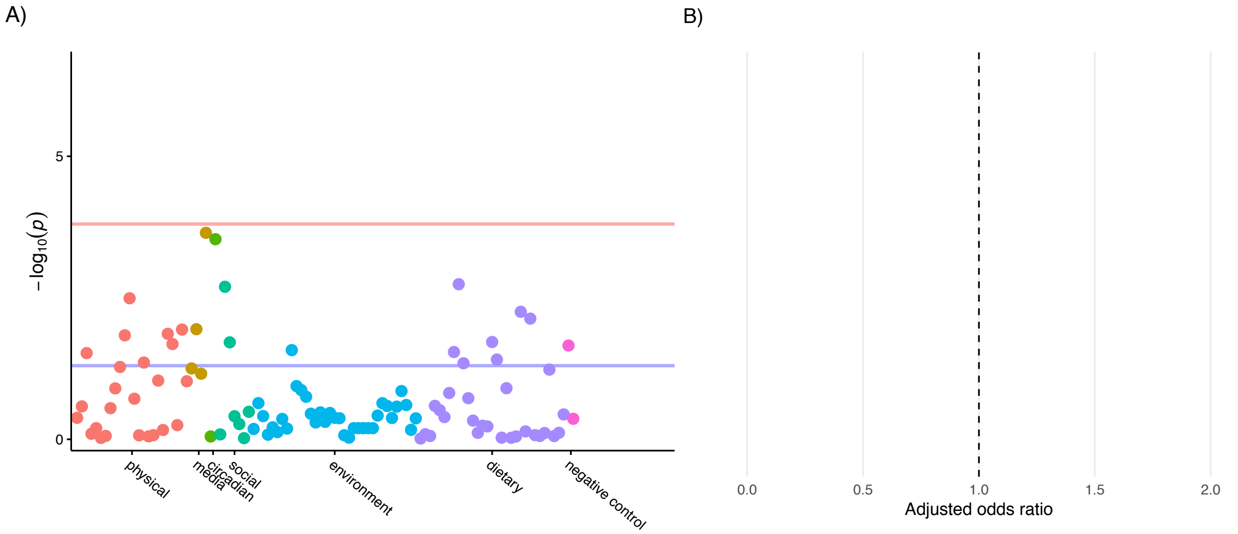

**Figure S22. Top hits for Model 2 (further adjusted for sociodemographic and health factors)**

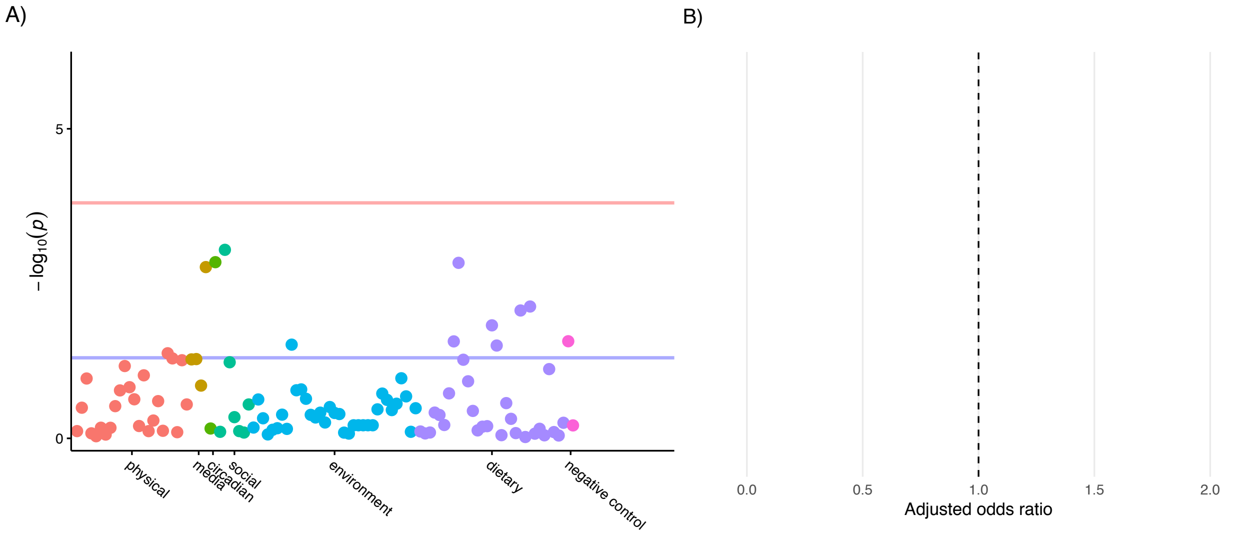

**Figure S23. All results for Model 0 (adjusted for base factors)**

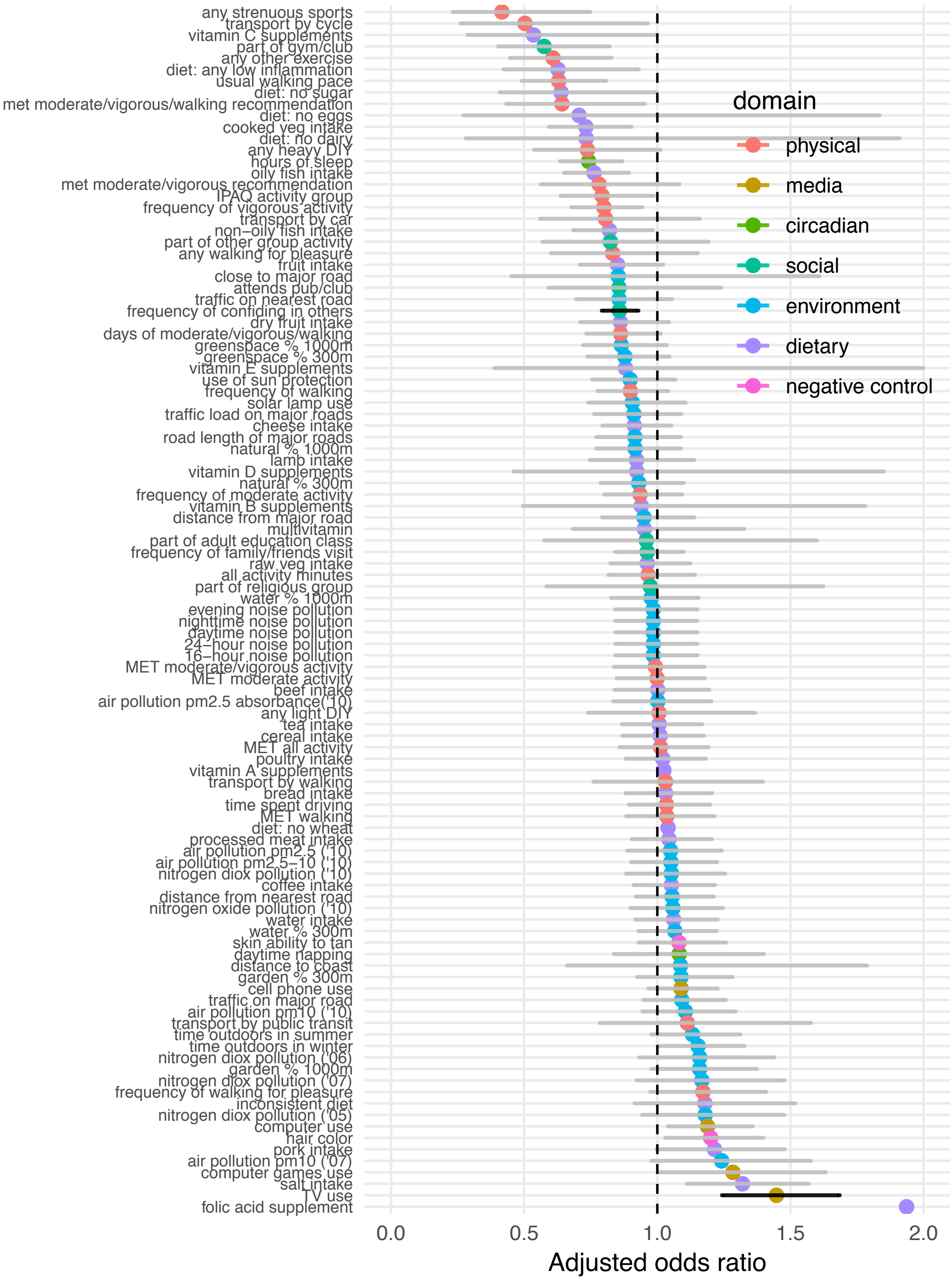

**Figure S24. All results for Model 1 (further adjusted for sociodemographic factors)**

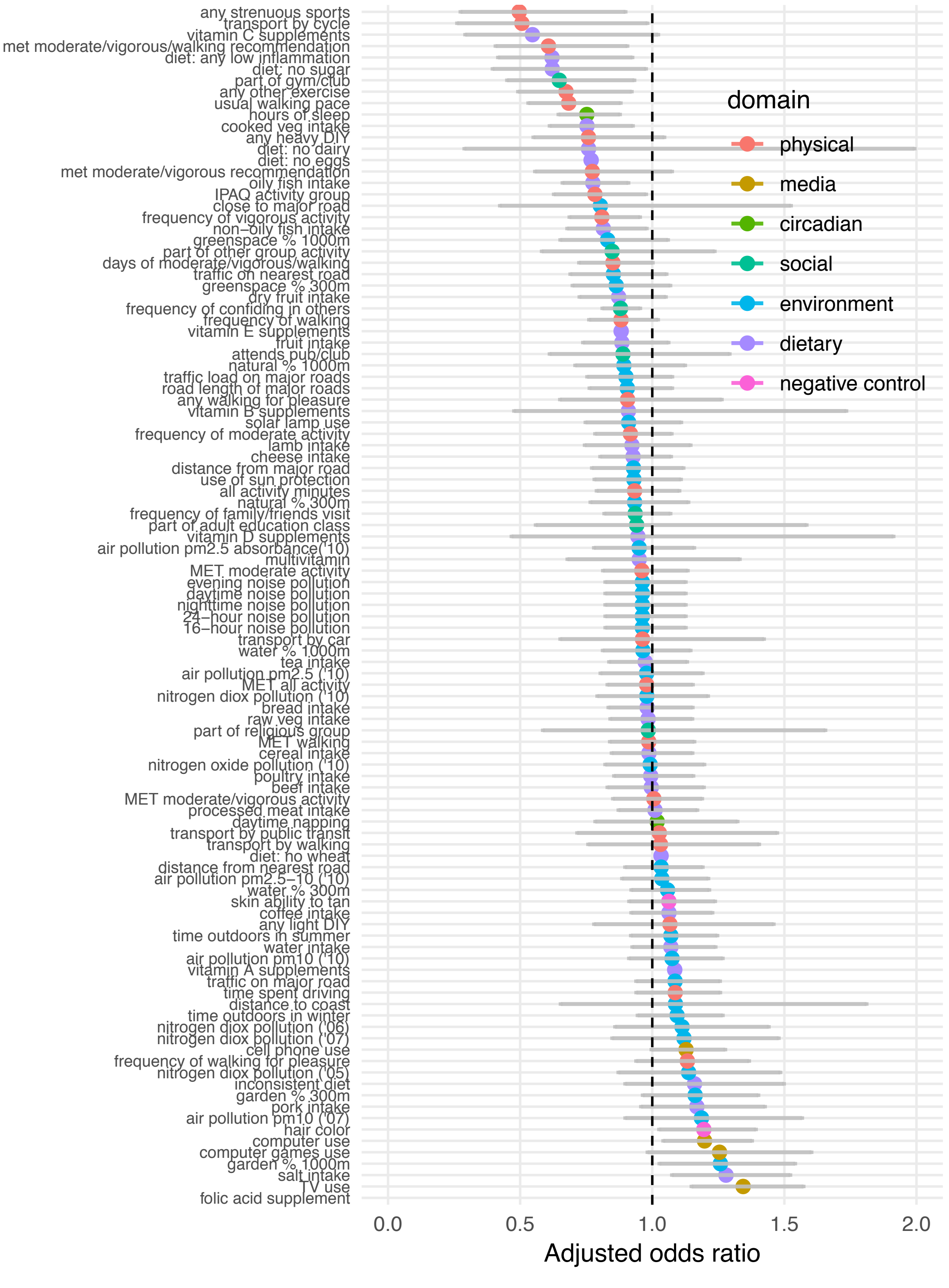

**Figure S25. All results for Model 2 (further adjusted for sociodemographic and health factors)**

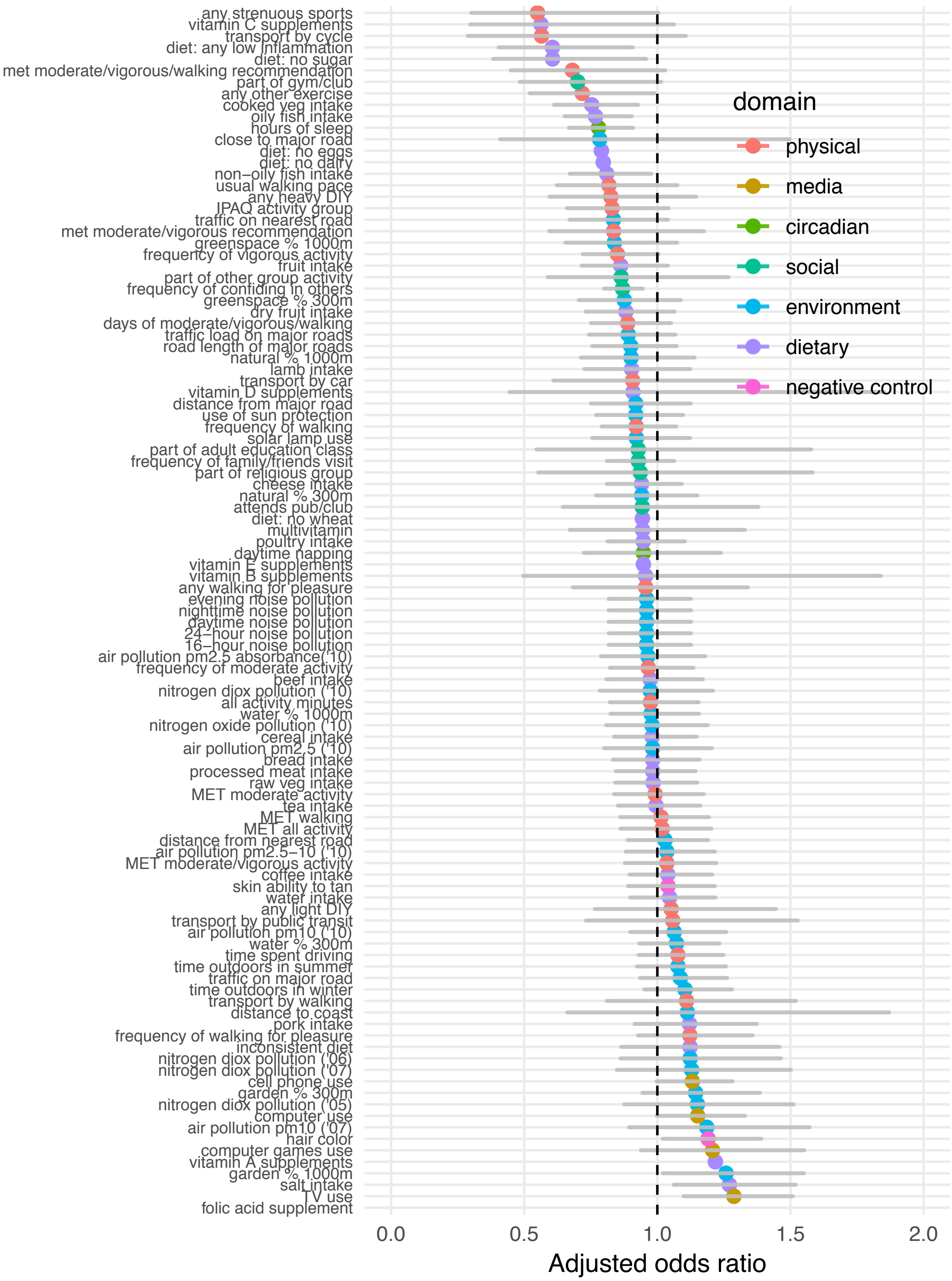

**Figure S26. MR estimates of top modifiable factors → the risk of depression with outliers removed, based on the weighted median method**

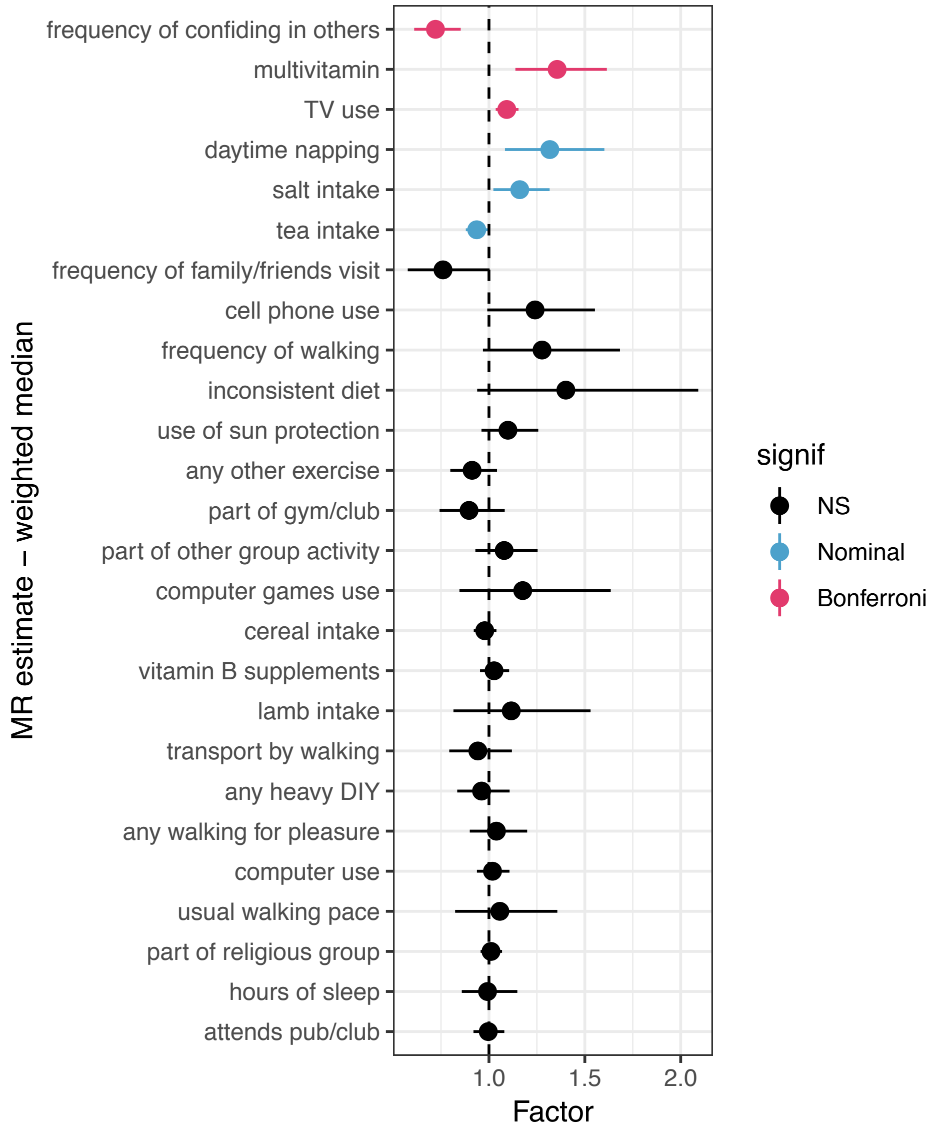

**Figure 27. MR estimates of depression → top modifiable factors with outliers removed, based on the weighted median method. Odds ratio estimates on left shown for dichotomous factors as outcomes, and beta estimates on right shown for non-dichotomous factors as outcomes.**

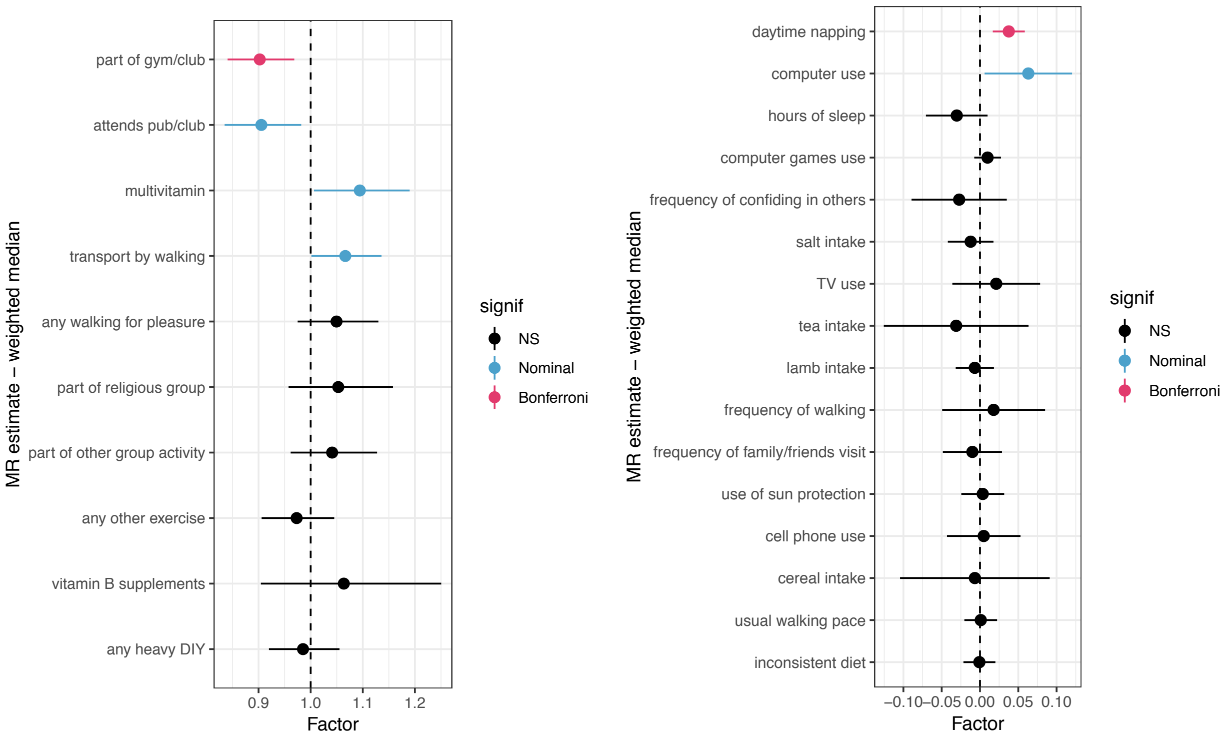
