## Supplementary Methods for "A two-stage approach to identifying and validating modifiable factors for the prevention of depression"

### Methods S0. Distinctions from previous UK Biobank studies of depression

A number of recent studies have examined depression in the UK Biobank, including a GWAS of various depression phenotypes (Howard et al., 2018), genome-wide gene-environment analyses of depression in the context of trauma exposure (Coleman et al., 2019, pre-print), and a phenome-wide association study examining relationships between depression PRS and a wide range of mental health/behavioral/imaging traits in the database, with accompanying Mendelian randomization analyses (Shen et al., 2019, pre-print). This study is distinct from previous work in notable ways, including its: (1) specific focus on identifying and testing *modifiable* factors associated with depression, rather than comorbidities and related traits/outcomes; (2) *prospective* examination of observational associations between modifiable factors and clinically significant depression among individuals who did not appear actively depressed at baseline; and (3) testing the effects of modifiable factors on depression *even among individuals at high polygenic risk*, rather than examining polygenic or genome-wide relationships of depression with other traits of interest. No study to our knowledge—in the UK Biobank or beyond—has reported these phenotypic and/or MR results to answer our set of prevention-oriented empirical questions.

**Methods S1. Genomic quality control**

We performed quality control on the genomic data based on metrics provided by the UK Biobank[23]. Specifically, we removed participants who were outliers for heterozygosity or data missingness, had putative sex chromosome aneuploidy, or were excluded from kinship inference. We also randomly removed one of each pair of subjects who were identified as third-degree relatives or closer. We retained participants with white British ancestry whose genomic data were used in autosome phasing and whose self-reported sex matched their genetically inferred sex. This resulted in an initial sample of 123,794 individuals of white British ancestry with high-quality genomic data in BGEN format.

**Methods S2. Further description of variables**

**A. Depression**

Baseline depression symptoms were measured via two Likert-scale items adapted from the PHQ-2 on depressed mood and anhedonia (response options ranging from “not at all” to “nearly every day”).

**B. Reported traumatic life events**

Exposure to childhood trauma was measured using items from a short form of the Childhood Trauma Questionnaire[27], of which three items pertaining to childhood physical, sexual, and emotional abuse were available. Each item was rated on a five-point Likert scale ranging from “never true” to “very often true.” Correspondingly, three items measuring exposure to partner-based physical, sexual, and emotional abuse, respectively, had been developed for the UK Biobank study on a similar severity scale as childhood trauma. Finally, four binary items assessed other lifetime traumatic events at any point prior to follow-up (i.e., exposure to sexual assault, physically violent crime, serious/life-threatening accident, and witnessing sudden violent death).

**C. Covariates**

We extracted baseline variables on participant characteristics (i.e., participant sex, age, assessment center), sociodemographic factors (i.e., socioeconomic deprivation, employment status, household income, completion of higher education, urbanicity, household size), and health factors (i.e., BMI, and physical illness/disability). Specifically, socioeconomic deprivation was indexed using the Townsend deprivation index, which was calculated based on the national census output area in which participants’ zip codes (at recruitment) are located. Household income was assessed average pre-tax total household income, with options ranging from <18K to

>100K. Individuals reported their educational qualifications and we derived a binary variable indicating the completion of higher education (i.e., college or university-level degree) as in previous research (Davies et al., 2016). Urbanicity was classified based on whether the participant was coded as living in an urban area of England/Wales or Scotland versus not, based on the home postal code matched to the 2001 census from the Office of National Statistics. Similarly, we derived a binary variable indicating current employment status, i.e., whether individuals endorsed paid employment or self-employment, and a binary variable indicating physical illness/disability, i.e., if individuals positively endorsed having any longstanding illness, disability, or infirmity. These variables were selected for inclusion as covariates as they could potentially bias the observed relationship between modifiable factors and depression (e.g., socioeconomic deprivation, urbanicity, or higher educational attainment may shape physical activity, diet, or media use patterns while also influencing depression risk), and were thus used to assess whether observed relationships between modifiable factors and depression would be robust to potential confounding in these domains. Here, we sought to include covariates previously linked to mental health that may not be readily modifiable.

**Methods S3. Data cleaning and processing**

We performed data cleaning on all variables and excluded variables that were missing for >20% of the sample. The dataset included multiple-choice categorical variables that were provided in array format based on the number of available choices (e.g., different types of activity) that we dummy coded into separate binary variables (i.e., yes/no for a given type of activity, assigning NA to those missing on the initial array variable). For continuous variables where a coding option of -10 indicated a value of less than 1 (e.g., less than one hour per day), we recoded this response to 0.5 to approximate less than 1 but greater than 0; substantive results were largely unchanged when recoding more conservatively to 0 (not shown). Some factor variables were recoded to reflect rational categories and/or a logical ordinal progression, where responses of increasing quantity were organized in ascending order. After these initial processing steps, negative values (e.g., -1 for "do not know", -3 for "prefer not to answer") indicated items that participants did not answer and were thus set as missing for all remaining variables, except the Townsend deprivation index which was scaled across negative and positive values.

**Methods S4. Polygenic scoring**

**Distribution of PRS-CS polygenic scores in the full analytic sample**

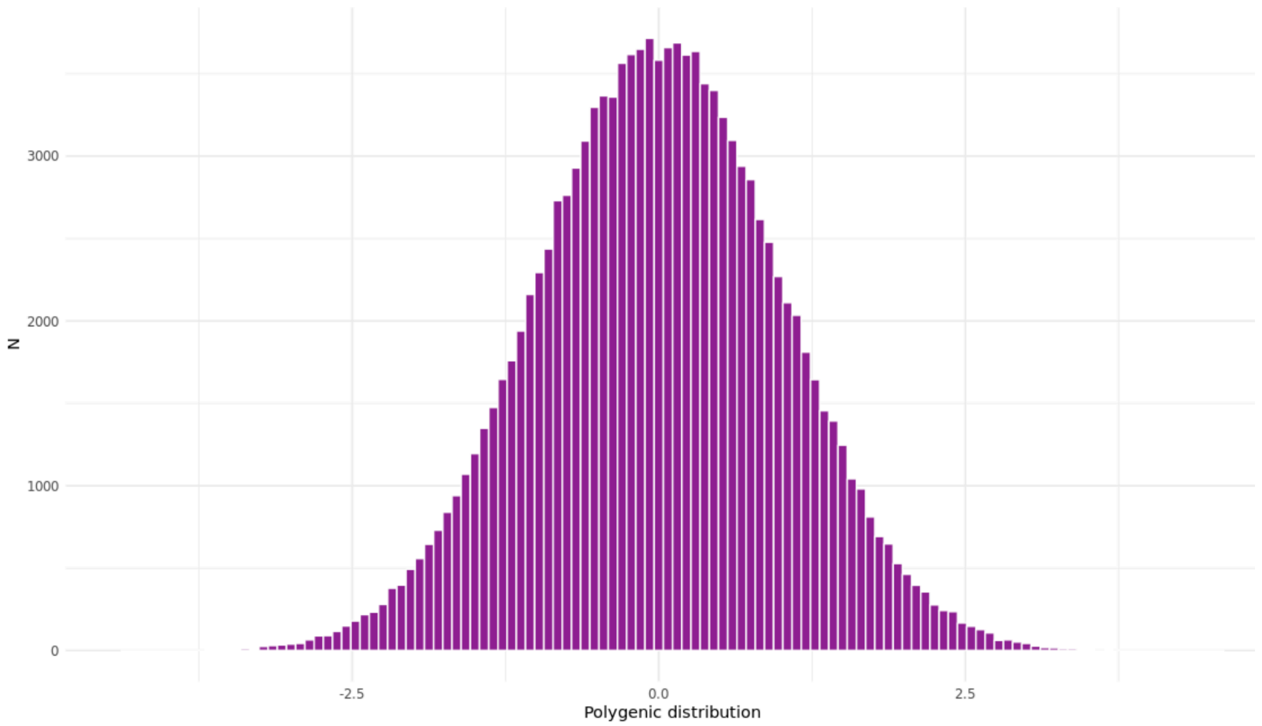


For comparison with the PRS-CS method (see distribution above and Methods in main manuscript), conventional clumping and thresholding procedures for polygenic risk scoring were performed using PRSice2 software. The below figures show PRSice2 estimates of explanatory $R^2$ of PRS across varying p-value thresholds in relation to follow-up depression, and well as distribution of PRS at the p-value threshold selected with highest explanatory $R^2$. Despite selecting the p-value threshold with highest explanatory $R^2$, the odds ratio associated with this conventional PRS (1.22) was slightly lower than for the PRS-CS (1.33) for follow-up depression.

**Explanatory variance of conventional PRS across p-value thresholds in relation to follow-up depression**

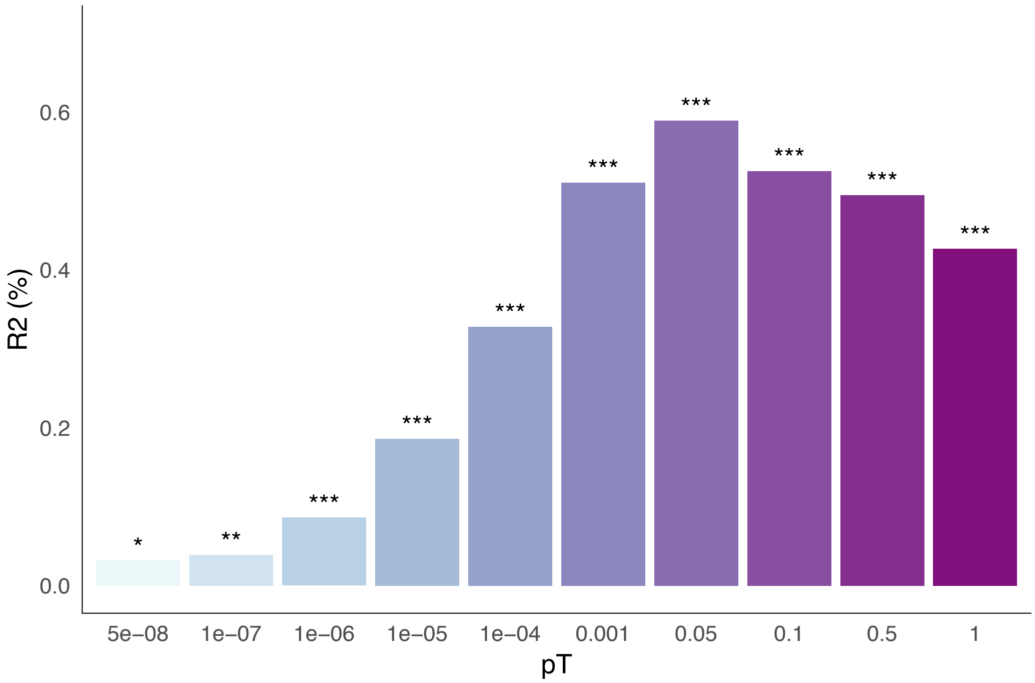


**Distribution of scores for the top conventional PRS (pT=0.05)**

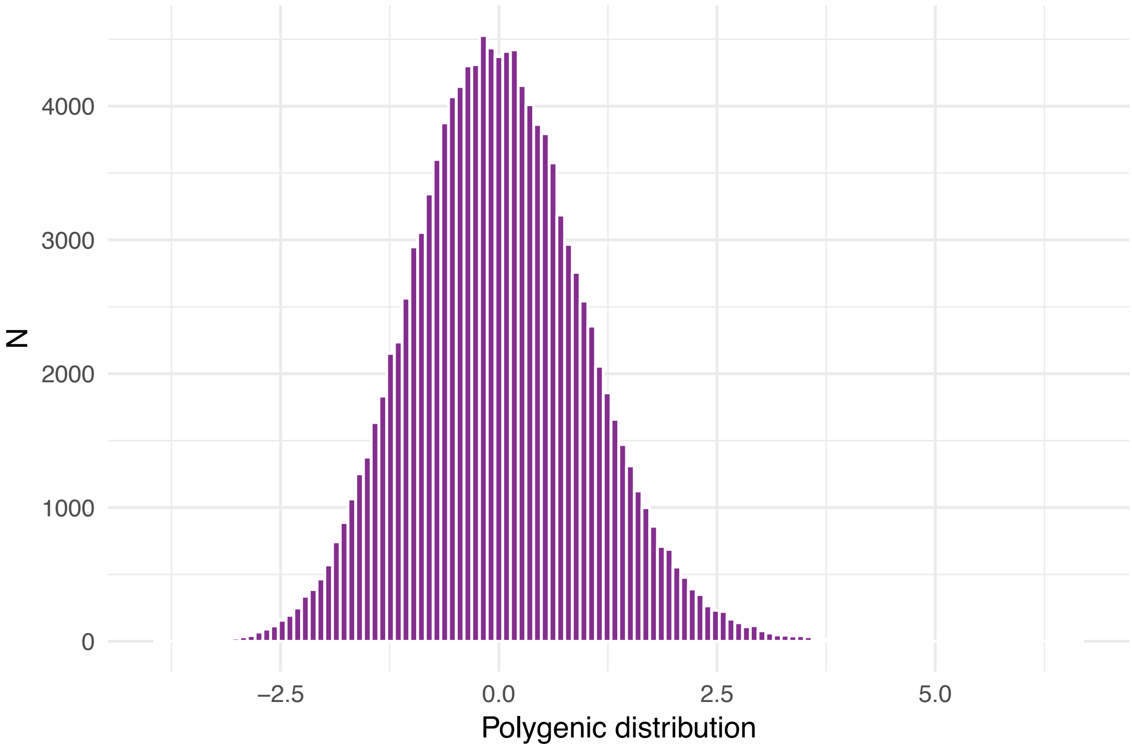

**Methods S5. Stratifying participants based on polygenic risk and reported traumatic life events**

As described in Methods (main manuscript), we randomly sampled a holdout training sample of 1,000 participants with an even split of cases and controls on follow-up depression. The rationale for a holdout sample of this size and case distribution was to derive predicted probabilities using the available risk factors in a smaller group of individuals drawn from the same study population for comparability, with standard enrichment of potential depression cases to improve statistical power for estimating the effects of each risk factor, while preserving relatively large sample sizes for the resulting analytic samples. We tested separate logistic regression models using clinically significant follow-up depression (binary variable) as the outcome, and (a) polygenic risk or (b) the set of reported traumatic life events, as independent variables. This strategy of estimating relative influences of different reported traumatic life events on follow-up depression based on a hold-out sample represented an alternative to the conventional strategy of assuming all events contribute equally to depression risk, which also has limitations. We used the resulting raw model coefficients derived from this training sample as variable weights to estimate predicted probability scores for follow-up depression among participants in the remaining testing sample (n=112,589) based on (a) polygenic risk or (b) the set of reported traumatic life events. The weights used for variables in each risk set are reported below, along with the distribution.

**Predicted probability scores for polygenic risk and stratification point**

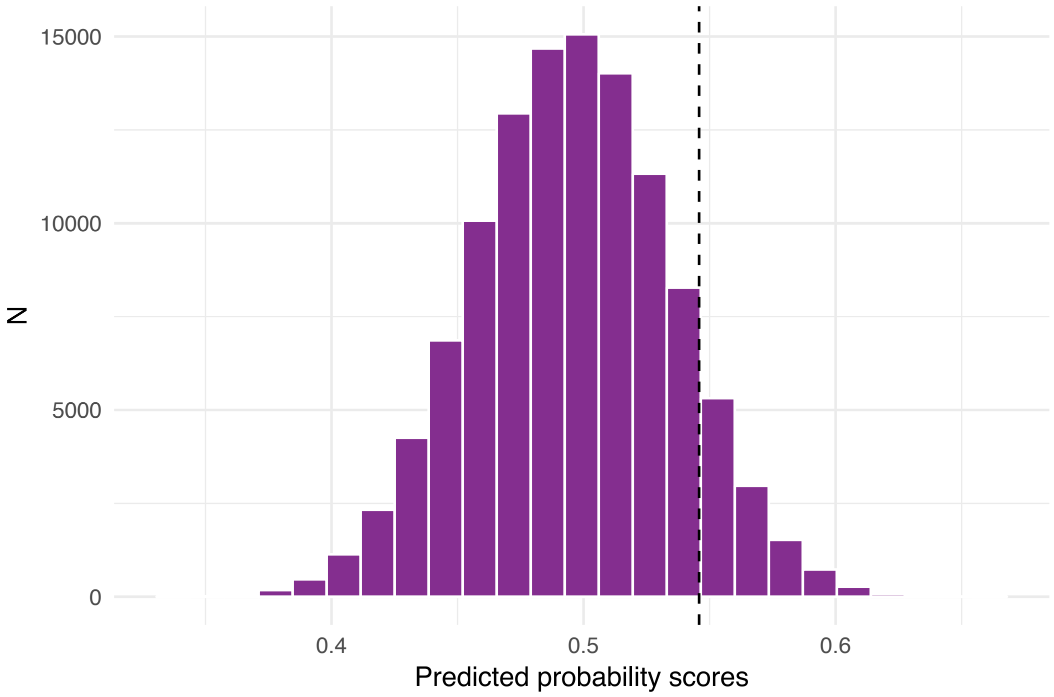


Predicted probability scores derived from training variable weights: Polygenic risk score (0.15867). Dashed line indicates cut-off point for establishing at-risk group (> 90th percentile).

**Predicted probability scores for traumatic life events and stratification point**

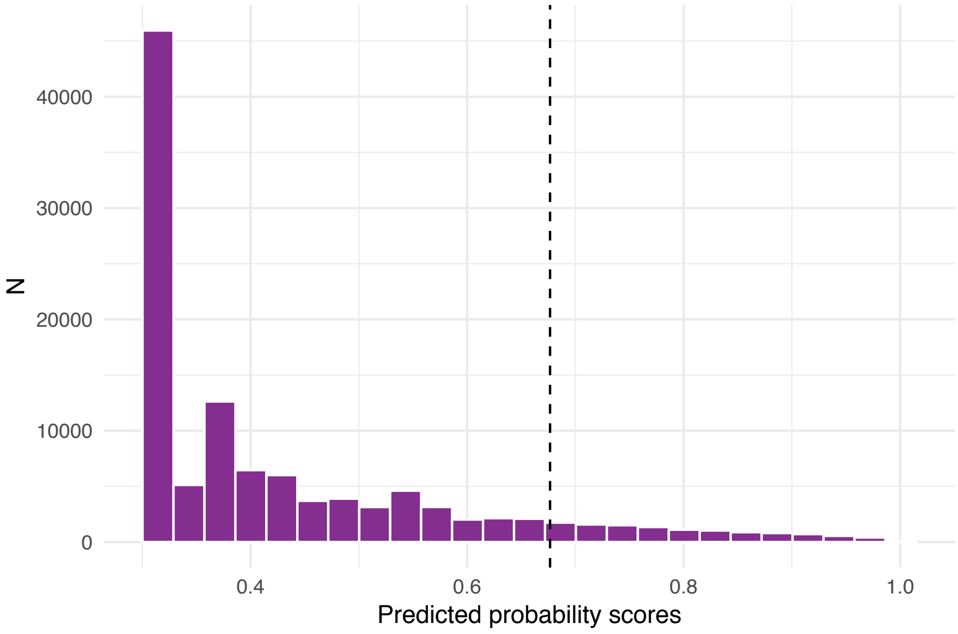


Predicted probability scores derived from training variable weights: Childhood physical abuse (0.10080), childhood emotional abuse (0.44014), childhood sexual abuse (0.11032), physical partner violence (0.07844), partner emotional abuse (0.48091), partner sexual interference (0.07409), lifetime exposure to sexual assault (0.31809), lifetime exposure to sudden violent crime (0.25048), life-threatening accident (0.44122), and witnessing sudden violent death (0.32587). Dashed line indicates cut-point for establishing at-risk group (> 90th percentile).

**Methods S6. Factors-wide association analyses**

Univariate associations between each baseline factor and follow-up depression status, adjusting for an increasingly stringent set of covariates as summarized in the main Methods, were tested using the *PheWAS* R package. Association tests between specific factors (as predictor) and depression (as outcome) were performed using all available data, using a logistic regression framework due to the binary nature of the depression phenotype. Dichotomous predictor variables were dummy coded for each model, while non-dichotomous (continuous or ordinal) variables were entered as predictors with linear assumptions. The analytic sample sizes for each tested association are summarized with all results in **Supplementary Tables 1a-i**.

**Methods S7. Two-sample Mendelian randomization analyses**

**A. Genetic instruments**

In a two-sample MR design, instruments can be extracted from summary statistics of large-scale, non-overlapping genome-wide association studies (GWAS). We accessed the GWAS Atlas online database[26] (https://atlas.ctglab.nl) to obtain publicly available summary statistics for each UK Biobank-based factor that was identified in the fully adjusted univariate association model for the full sample. GWAS Atlas summary statistics that were missing for five UK Biobank derived variables (i.e., meeting recommendations for moderate/vigorous/walking activity) were not tested in MR framework; however, related traits (e.g., walking frequency) were examined as possible. For depression, we retained the same set of summary statistics[12] for major depression used in previous steps for polygenic scoring in this sample. We clumped SNPs correlated at $r^2 > .0001$ in a 1000 kb window for independence based on European ancestry reference data from the 1000 Genomes Project. For genetic instruments, we selected highly associated SNPs (discovery GWAS p-value < 1x10e-7) for the exposure of interest.

**B. Additional details for MR analyses**

Using the *TwoSampleMR* package in R, we conducted MR analyses to estimate the effect of each modifiable factor on major depression, and vice versa. The *TwoSampleMR* package harmonizes exposure and outcome datasets containing information on SNPs, alleles, effect sizes (odds ratios converted to betas via log transformation), standard errors, and p-values. In this study, ambiguous SNPs could not be inferred due to missing effect allele frequency information in GWAS Atlas summary statistics, and were therefore removed for analysis. Prior work has suggested that inclusion/exclusion of ambiguous SNPs does not substantively change MR results.

**C. Nominal MR results**

Other findings were nominally significant at the traditional $p<0.05$ threshold, which suggested potential beneficial effects of tea intake (OR=0.95, 95% CI [0.91-0.99], p=1.63E-2); more frequent social visits (OR=0.79, 95% CI [0.65-0.96], p=1.80E-2; non-significant WM estimate though directionally consistent) and engaging in exercises like swimming and cycling (OR=0.89, 95% CI [0.81-0.98], p=1.95E-2; non-significant WM estimate though directionally consistent), as well as deleterious effects of salt intake (OR=1.10, 95% CI [1.01,1.19], p=3.45E-2) on the risk of depression. We also observed nominal evidence in the reverse direction, suggesting depression may be associated with reduced gym/club use (OR=0.93, 95% CI [0.88-0.98], p=7.10E-3), attendance at pubs/social clubs (OR=0.92, 95% CI [0.87-0.98], p=8.03E-3), and time spent driving (beta=-0.05, 95% CI [-0.09, -0.015], p=6.67E-3), as well as increased computer use (beta=0.04, 95% CI [0.14-0.09], p=7.97E-3) and playing computer games (beta=0.01, 95% CI [0.001-0.03], p=3.27E-2; non-significant WM estimate and also directionally inconsistent), and supplementation not only with multivitamins (as noted earlier) but also vitamin B (OR=1.14, 95% CI [1.02-1.27], p=1.89E-2; non-significant WM estimate though directionally consistent). Although walking was phenotypically associated with reduced odds for depression, MR results suggested that depression may be nominally associated with increased tendency for walking, whether for pleasure or as a form of transportation (OR=1.05, 95% CI [1.004-1.11], p=3.61E-2, non-significant WM estimate though directionally consistent; OR=1.06, 95% CI [1.009-1.11], p=1.87E-2, respectively).
